# R-Loops between nascent pri-miRNAs and the encoding loci promote co-transcriptional processing of miRNAs in plants

**DOI:** 10.1101/2021.09.07.459282

**Authors:** Lucia Gonzalo, Ileana Tossolini, Tomasz Gulanicz, Damian A. Cambiagno, Anna Kasprowicz-Maluski, Jan Dariusz Smolinski, Sebastian Marquardt, Zofia Szweykowska-Kulinska, Artur Jarmolowski, Pablo A. Manavella

**Author notes:** These authors contributed equally (Lucia Gonzalo, Ileana Tossolini, Tomasz Gulanicz).

## Abstract

In most organisms, the maturation of nascent RNAs is coupled to transcription, undergoing many processing steps co-transcriptionally. Unlike in animals, the RNA polymerase II (RNAPII) transcribes microRNAs (miRNAs) as long and structurally variable pri-miRNAs in plants. Current evidence suggests that the miRNA biogenesis complex assembly initiates early during the transcription of pri-miRNAs in plants. However, it is unknown whether miRNA processing occurs co-transcriptionally. Here, we show that plant miRNA biogenesis is coupled to transcription in a process that relies on the formation of DNA:RNA hybrids (R-loops) between the nascent transcript and the encoding loci. We used native elongating transcript sequencing data and imaging techniques to demonstrate that plant miRNA biogenesis occurs co-transcriptionally. We found that the entire biogenesis occurs coupled to transcription for pri-miRNAs processed from the loop but requires a second nucleoplasmic step for those processed from the base of the hairpin. Furthermore, we found that co- and post-transcriptional miRNA processing mechanisms co-exist for most miRNAs in a dynamic balance. Notably, we discovered that R-loops between the 5’-end single-stranded arm of the pri-miRNAs and the encoding loci anchor the transcript, promoting co-transcriptional processing. Our data demonstrate the coupling of transcription and miRNA processing in plants and discovered an unexpected function for R-loops promoting RNA processing. Furthermore, our results suggest the neo-functionalization of co-transcriptionally processed miRNAs, boosting countless regulatory scenarios.

## Introduction

The biogenesis of miRNAs in plants is a unique and evolutionary divergent pathway that differs from its counterpart in metazoans (Axtell et al., 2011; Zhang et al., 2018). In plants, for example, independent transcriptional units containing specific promoters, terminator and even introns encode most miRNAs (Cuperus et al., 2011; Szarzynska et al., 2009). The RNA polymerase II (RNAPII) transcribes plant *MIRNA* loci as capped and polyadenylated primary transcripts (pri-miRNAs). Unlike the animal pathway, where DROSHA and DICER concatenate to produce the mature miRNAs, a single processing complex containing DICER-Like 1 (DCL1) conducts the entire process inside the plant nucleus (Achkar et al., 2016; Bartel, 2018).

Opposite to pri-miRNAs in metazoans, which are homogeneous in size, plant’s pri-miRNAs are highly variable in length and secondary structure ranging from hundreds to thousands of base pairs in polycistronic transcripts (Bologna et al., 2013; Moro et al., 2018; Singh et al., 2020). This particularity confronts DCL1 with a problem: recognizing the position of the active miRNA within such variable pri-miRNAs. Consequently, the miRNA-processing complex of plants relies on accessory proteins, such as HYPONASTIC LEAVES1 (HYL1) and SERRATE (SE), and structural features in the pri-miRNAs to guide DCL1 to the precise slicing positions {Dong, 2008 #13;Re, 2019 #14;Rojas, 2020 #120;Yang, 2014 #16}. As a result, alternative processing modes take place depending on the characteristics of each pri-miRNA (Moro *et al*., 2018; Zhu et al., 2013). In most cases, DCL1 produces a first cut near the base of the hairpin structure in a process known as Base-to-Loop processing (BTL) that resembles the processing from pri- to pre-miRNA by Drosha in animals (Moro *et al*., 2018). In other cases, the processing complex recognizes features in the terminal loop and initiates DCL1-mediated processing from the hairpin loop to the base (LTB) (Addo-Quaye et al., 2009; Bologna et al., 2009; Bologna *et al*., 2013; Moro *et al*., 2018; Zhu *et al*., 2013). Interestingly, sequential cuts of the pri-miRNA every ∼21 nt by DCL1 releases the mature miRNA from long precursors (BTLs and LTBs) (Bologna *et al*., 2013; Moro *et al*., 2018).

Some components of the miRNA-biogenesis complex, such as DCL1 and HYL1, are located in nuclear speckles known as dicing bodies (D-Bodies) but alto associated with the *MIRNA* loci (Bhat et al., 2020; Cambiagno et al., 2021; Fang et al., 2015; Kim et al., 2011; Wang et al., 2013). The recruitment of DCL1 to the *MIRNA* loci relies on its association with the RNAPII-accessory complexes MEDIATOR and ELONGATOR. In the first case, HASTY (HST), the plant ortholog of human EXPORTIN 5, acts as a scaffold stabilizing the interaction of MED37-DCL1 that allows the recruitment of DCL1 to nascent pri-miRNAs (Cambiagno *et al*., 2021). Similarly, DCL1 recruitment to the *MIRNA* relies on the elongator complex (Fang *et al*., 2015). Such recruitment of the processing machinery to *MIRNA* loci may suggest that miRNA biogenesis takes place co-transcriptionally in plants, as described for animals (Ballarino et al., 2009; Morlando et al., 2008; Nojima et al., 2015; Pawlicki and Steitz, 2008). However, opposite to co-transcriptional splicing, which occurs progressively as the mRNA is transcribed, the co-transcriptional processing of plant pri-miRNAs would first need the transcription and folding of the entire step-loop region before it can be recognized and processed. This particularity gives a small temporal window for the processing to occur co-transcriptionally before the pri-miRNA is released to the nucleoplasm, and would likely involve some sort of RNA anchoring. Thus, it opens the question of whether the recruitment of the processing complex to the *MIRNA* loci induces co-transcriptional processing or only allows an earlier complex assembly. Therefore, determining whether the plant pri-miRNAs can be co-transcriptional processed is one of the most outstanding open questions in the field.

Co-transcriptional RNA processing is frequent in most organisms (Bentley, 2014; Herzel et al., 2017; Lee and Tarn, 2013; Peck et al., 2019; Yang et al., 2021). Such processes commonly co-exist with a post-transcriptional processing counterpart, and the balance between them can be regulated to produce alternative physiological outcomes. DNA:RNA hybrid (R-Loop) formation is also a common co-transcriptional event (Chedin, 2016; Hamperl and Cimprich, 2014). The R-loops are naturally occurring DNA-RNA hybrids formed either in *cis,* with the RNA encoding locus, or in *trans* due to sequence complementarity. *Cis* R-loops commonly involve the nascent transcript, especially during slow transcription (Chedin, 2016; Zatreanu et al., 2019). These chromosomal structures are frequent in bacteria, yeast, animals, and plants, playing roles in many biological processes (Crossley et al., 2019; Ginno et al., 2012; Niehrs and Luke, 2020; Santos-Pereira and Aguilera, 2015; Xu et al., 2017). In plants, R-loops play roles in development, gene regulation, and genome integrity (Ariel et al., 2020; Shafiq et al., 2017; Sun et al., 2013; Yang et al., 2017; Yang et al., 2020; Yuan et al., 2019).

Here, we used plant native elongating transcripts sequencing (plaNET-seq) data to profile genome-wide nascent pri-miRNA processing intermediates associated with the RNAPII. The results indicated that pri-miRNAs are processed co-transcriptionally in Arabidopsis. This was also confirmed using different microscopic approaches of nascent pri-miRNAs. Furthermore, we found that once initiated, co-transcriptional processing can occur entirely associated with the transcriptional complex, in the case of LTB and LTBs miRNAs, or in a two-stages fashion, resembling animal’s pri- to pre-miRNA processing, for miRNAs processed from the base. We also discovered that co-transcriptional and post-transcriptional processing co-exist and fluctuate between growth conditions for most pri-miRNAs. Surprisingly, we found that co-transcriptional processing of pri-miRNAs largely relies on R-loops between the nascent transcripts and the *MIRNA* encoding loci to stabilize the precursor and allows DCL1 processing. Finally, our data suggest that regulation of R-loops formation directly impacts on whether a pri-miRNA is processed co-transcriptionally.

Overall, our study identified an alternative miRNA biogenesis pathway, discovered an unexpected function for R-loops promoting RNA processing, and opened the doors to neo-functionalization of co-transcriptionally processed miRNAs with the concomitant regulatory implications.

## RESULTS

### Imaging of pri-miRNAs suggest co-transcriptional miRNA biogenesis

Current knowledge regarding the assembly of the miRNA-biogenesis complex suggests that it is possible that the processing of miRNAs is linked to transcription. To investigate whether miRNA-biogenesis and transcription are coupled, we first used Fluorescence *in situ* Hybridization (FISH) to visualize pri-miRNAs within the nucleus using confocal microscopy. For these experiments, we selected pri-miR163 and pri-miR156a as both contain introns that allow us to differentiate nascent pri-miRNAs from the mature molecules (Figure S1A). We designed the FISH probes to target an intron or exon located downstream of the stem-loop (probes named Intron and Exon, respectively), the spliced pri-miRNA transcript (Exon/Exon), the loop region (Loop), the mature miRNA region (miRNA), or the miRNA complementary sequence (miRNA*) (Figure S1A). The results showed that pri-miRNA156a and pri-miRNA163 localized in one or two discrete fluorescence spots within the nucleoplasm (Figure 1A and S1B). We then validated these results using the Stellaris FISH RNA method to detect pri-miRNA156a in nuclei of *A. thaliana* cells. The probes were designed against the intron sequence of pri-miRNA156a and labelled with either Quasar 570 or fluorescein (6-FAM). Again, these experiments showed that pri-miRNA156a accumulated in one or two well-defined nuclear spots (Figure S1C).

**Figure 1.**
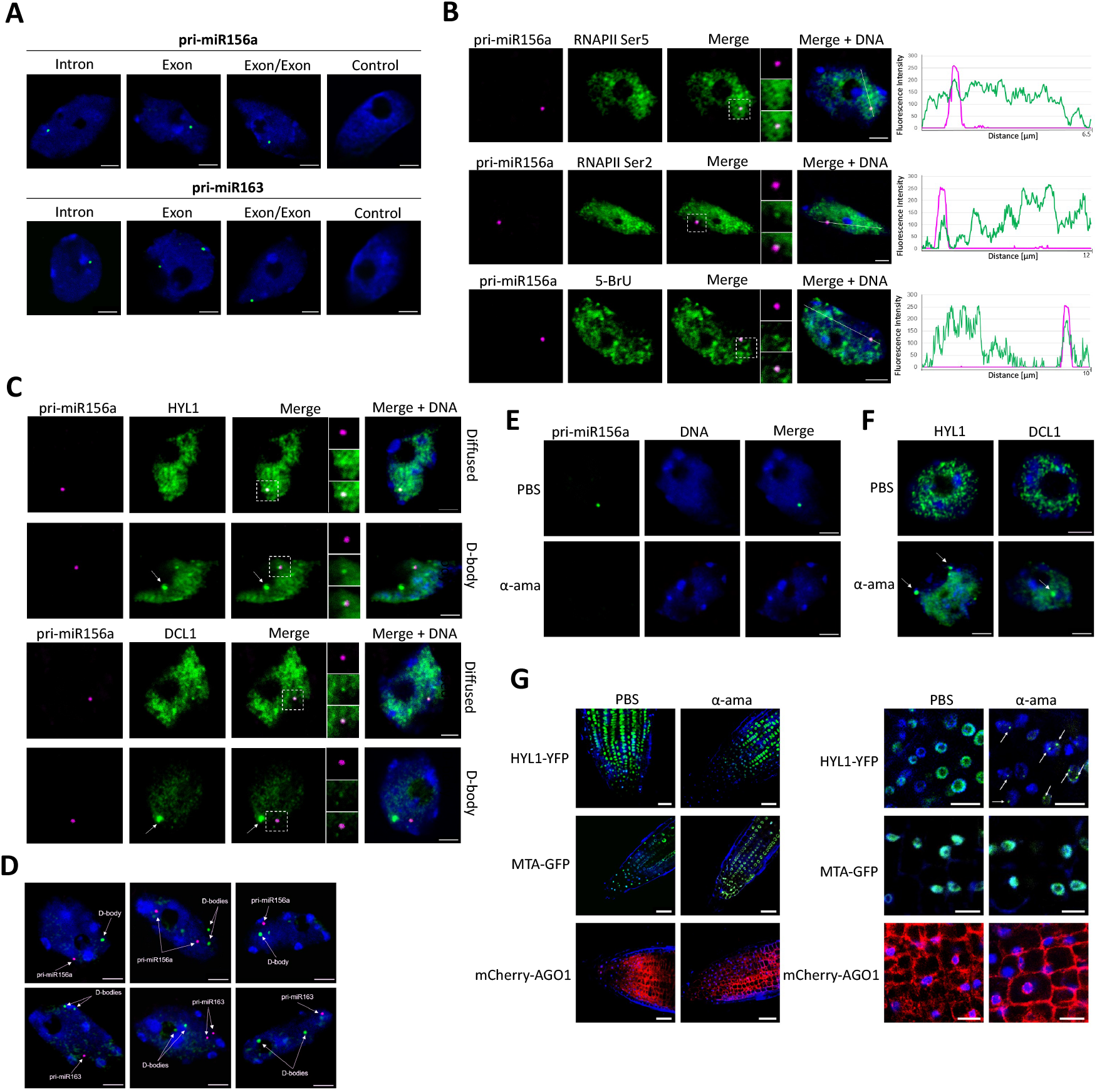
Subcellular distribution and interactions of nascent pri-miRNAs and the miRNA biogenesis complex. (A) FISH of pri-miRNA156a and pri-miR163 (green) using mouse antibodies targeting digoxigenin in the nuclei isolated from wild-type plant cells. The probes hybridizing to intron (Intron), exon (Exon), transcripts after splicing (Exon/Exon); hybridization buffer without any probe was used as a negative control (Control). (B) Detection of pri-miR156a using RNA Stellaris probes (magenta) and RNAPII Ser5, RNAPII Ser2, or newly synthesized transcripts (green). On the right are shown the fluorescence intensity plotted along the white line shown each panel on the left. The fluorescence intensity of pri-miR156a is depicted as the magenta curves, while RNAPII Ser 5, RNAPII Ser 2, and 5-BrU as the green curves. (C and D) FISH of pri-miRNA156a (magenta) combined with immunolabeling of HYL1 (C) or DCL1 (C and D). (E and F) FISH of pri-miRNA156a using RNA Stellaris (green) (E) and immunodetection of HYL1 or DCL-1 (green) (F) in plants treated with α-amanitin or PBS as a negative control. White arrows mark D-Bodies. In all cases, the nuclei were stained with Hoechst (blue). Scale bar - 2.5 μm. (G) HYL1-YFP distribution in the root cells of the meristematic zone in plants treated with buffer (PBS) (upper panel) and α-amanitin (bottom panel). In the right is the magnification of representative images obtained from the root meristematic zone cells of four different plants. White arrows mark D-Bodies. Scale bar: 10 µm. DNA was stained with Hoechst (blue).

The localization of these pri-miRNA transcripts in one/two discrete loci may perfectly reflect the transcription sites of both copies of each gene. To confirm it, we used RNA Stellaris probes to analyzed subnuclear localization of pri-miRNA156a side by side with RNAPII immunodetection, using antibodies specific to the C-terminal domain (CTD) serine 5 and serine 2 isoforms (RNAPII^Ser5^, RNAPII^Ser2^). We also apply the 5-BrU incorporation method (Niedojadlo et al., 2012) to visualize newly formed transcripts. Our results showed that pri-miRNA156a co-localized with both transcriptionally active RNAPII and 5-BrU (Figure 1B). These results confirm that the detected spots are the pri-miRNAs transcription sites. Interestingly, we did not detect additional spots, even when using probes that could detect post-transcriptionally processed pri-miRNAs. Such distribution was also observed in animal cells by different microscopy approaches (Salataj et al., 2019; Turunen et al., 2019). This could probably represent either a very quick nucleoplasmic processing of pri-miRNAs or a diffused processing, which in turn challenges the role of D-Bodies during miRNA processing. These results encouraged us to test whether pri-miRNAs also co-localize with the miRNA biogenesis complex. Thus, we visualized pri-miRNA156a using RNA Stellaris probes followed by immunolocalization of microprocessor proteins HYL1 and DCL1. We observed pri-miR156a in one or two discrete foci while both HYL1 and DCL1 localized either dispersed in the nucleoplasm or in well-defined nuclear bodies, the so-called D-Bodies (Figure 1C and S1D). This dual distribution of HYL1 and DCL1 coincides with previous reports (Song et al., 2007). We found both proteins predominantly distributed in the nucleoplasm (∼70% of all tested cells), whereas DCL1- or HYL1-containing nuclear bodies were observed in roughly 30% of cells (Figure S1D). Coincident with the reports locating DCL1 and HYL1 in *MIRNA* loci (Bhat *et al*., 2020; Cambiagno *et al*., 2021; Fang *et al*., 2015), our results showed that these two proteins co-localized with pri-miR156a transcript but not into the D-Bodies (Figure 1C, and S1E). We repeated these experiments including pri-miR163, and adjusting the stringency and acquisition parameters to focus only on well-defined structures (D-Bodies and transcription sites). Again, we observed that both pri-miRNAs transcription sites did not match D-Bodies (Figure 1D).

Altogether, our microscopy results support the idea of co-transcriptional miRNA processing, as the complex is assembled on pri-miRNA transcription sites. They also indicated that the processing complex contained in D-Bodies is not associated with the transcriptional complex. Whether these nuclear structures represent post-transcriptional pri-miRNA processing places or they are simply reservoirs of inactive proteins remain to be addressed. The relatively low number of cells displaying D-Bodies may also play against an active function in miRNA biogenesis. To further explore this aspect, we treated cells with α-amanitin to inhibit RNAPII activity and we repeated the pri-miRNA FISH and HYL1/DCL1 immunostaining. As expected, we did not register any fluorescence signal of pri-miRNA156a in plants treated with α-amanitin, confirming that transcription was successfully blocked (Figure 1E). Notoriously, we observed a shift in HYL and DCL1 subnuclear localization upon RNAPII inhibition by α-amanitin towards accumulation in nuclear bodies (Figure 1F). We confirmed this observation *in planta* by analyzing HYL1-YFP distribution in roots of 10-day-old *A. thaliana* plants treated with α-amanitin. After 2 hours of incubation, we detected changes in the localization of the protein toward nuclear bodies containing HYL1-YFP (Figure 1G and S1F). In contrast, we did not observe any changes in the distribution of two other fluorescent fusion proteins, MTA-GFP and mCherry-AGO1, used as controls after α-amanitin treatment (Figure 1G). These results indicate that RNAPII inhibition increased the number of root meristem cells containing D-Bodies, supporting a scenario where D-Bodies are not the primary source of miRNA processing acting perhaps as reservoir of miRNA biogenesis proteins.

### Plant pri-miRNAs are processed co-transcriptionally

The numerous reports describing the association of miRNA biogenesis factors with pri-miRNAs encoding loci and our microscopy data prompted us to test whether such recruitment triggers co-transcriptional processing of miRNAs. To test this hypothesis, we first immunoprecipitated (IP) nascent transcripts using an antibody against the RNAPII (RIP) followed by the detection of processing intermediates by modified 5’-RACE. Plants expressing a HIS-tagged version of the ATHB1 transcription factor (Miguel et al., 2020) and an anti-HIS antibody were used as a negative control for RIP-5’-RACE. We detected processed fragments corresponding to the reported DCL1-mediated cleavage site associated with the RNAPII, but not to AtHB1, for the three tested pri-miRNAs (Figure 2A, S2A). For the pri-miR156a, where processing proceeds from the loop to the base, we detected both processing intermediates associated with the RNAPII (Figure 2A, S2A). Aiming to confirm this observation at a genome-wide scale, we used plaNET-seq data (Kindgren et al., 2020; Leng et al., 2020) to identify pri-miRNA processing intermediates produced from nascent transcripts. In plaNET-seq, a FLAG-tagged version of NRPB2, transformed into a *nrpb2*-1 mutant background, is immunoprecipitated and nascent transcripts associated with the RNAPII are detected by Illumina sequencing (Kindgren *et al*., 2020). We first aligned the processed reads, from wild-type Col-0 plants, to the *MIRNA* loci in the Arabidopsis genome. The 5’ nucleotide (nt) of each read was then plotted to identify co-transcriptional processing intermediates. *MIRNA* loci were scaled from miRNA-5p to the miRNA-3p to a fixed length to visualize the general processing profile. In addition, pri-miRNAs were sorted depending on their processing direction (BTL, LTB and their sequentially processed counterparts BTLs and LTBs), (Moro *et al*., 2018). Co-transcriptional processing intermediates, visualized as a peak in the nucleotide right after the known DCL1 cleavage sites, were easily detected, showing a clear global pattern (Figure 2B, cyan arrow). Pri-miRNA transcription start sites (TSS) were observed as a region rich in undefined peaks, as expected from the variable-length from TSS to the scaled region for each individual pri-miRNA (Figure 2B, orange mark). We observed a similar narrow peak when we plotted the data to the un-scaled miRNA-3p region but not the miRNA-5p region, as expected from the processing direction (Figure 2C).

**Figure 2.**
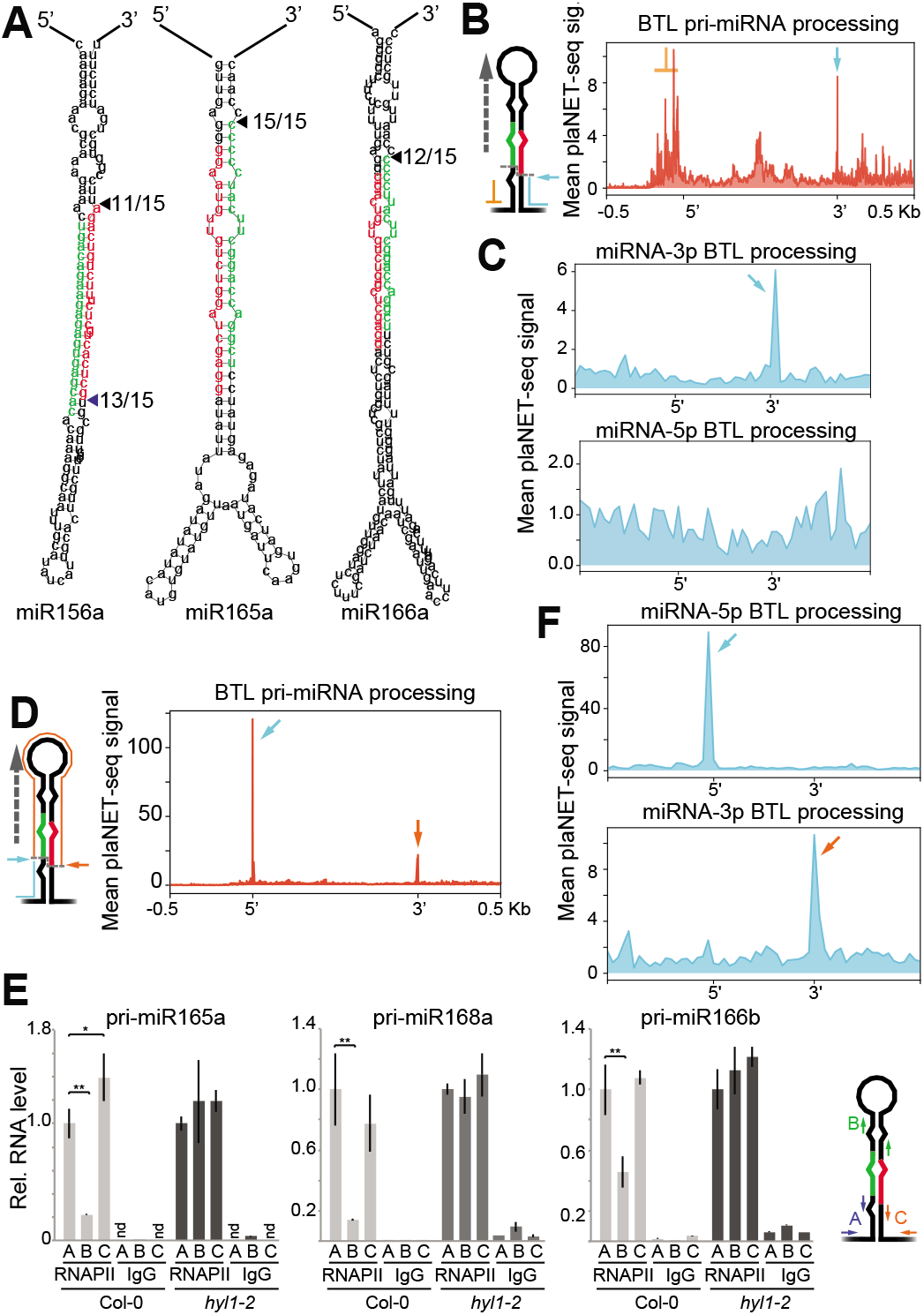
Plant pri-miRNAs are processed co-transcriptionally. (A) Detection of nascent pri-miRNA processing intermediates associated with RNAPII as detected by 5’RACE. The fraction of clones with the expected 5’ end (among all the sequenced products) is indicated next to the predicted pri-miRNAs secondary structures. The miRNA-5p and miRNA-3p positions are noted in red and green respectively. (B, C, D, and F) Metagene analysis of nascent BTL pri-miRNAs processing intermediated associated to RNAPII as determined by scoring the 3’-end nucleotide in of plaNET-seq reads in the plus and minus strands of plaNETseq samples (B and D) or plaNETseq negative controls (C). Pri-miRNAs were scaled from the beginning of miRNA-5p to the end of miRNA-3p (B and C) or using only the mature miRNAs sequences (D). Cyan and orange arrows indicated the processing fragments detected also marked in Figure 1. A scale identical to the plot displayed in Figure 2D is shown in the left panel of (C) while a zoom in is shown in the right panel. (E) Co-transcriptional pri-miRNA processing score as measured by RT-qPCR in RNAPII or IgG IP RNA-samples from Col-0, and hyl1-2. Error bars corresponds to 2xSEM. P<0.05 (*) or <0.01 (**), in a two-tailed unpaired T-Test, were considered statistically significant. RT-qPCR using primers pairs A, B and C in the IP fraction were normalized to U6 transcript and to the input levels using the same primers. Co-transcriptinal processing was measured as a reduction in the hairpin amplicon (primers B) relative to 5’ region (amplicon A). Non-detected RNA levels are displayed as n.d.

Interestingly, when we plotted the 3’-nt of the plaNET-seq mapped reads, but not from the negative control, we detected the processing intermediates with even cleaner profiles (Figure 2D, S2B, S2C). The detection of such peaks suggested the retention of the 5’-end of pri-miRNAs in the transcriptional complex after processing. Such a scenario is observed in co-transcriptionally spliced mRNAs where processed exons remain bound to the complex until the splicing finish (Nojima *et al*., 2015). Considering that analyzing the 3’-nt end of the mapped reads eliminates the noise of TSSs and fragmented molecules, and provides cleaner profiles, we used this approach from now on. To further confirm this result, we performed a RIP experiment using an H3 antibody to evaluate pri-miRNA processing fragments associated with the chromatin. Confirming the previous observation, we found both the 3’- and 5’-ends pri-miRNAs processing-fragments associated with the chromatin, but a relative depletion of the stem-loop processed regions (Figure 2E). Such reduction of the hairpin region, which is not evident in the processing-defective mutants *hyl1-2*, is compatible with co-transcriptional processing of the pri-miRNAs (Figure 2E). The surprising retention of the 5’ processed region of the pri-miRNAs in the transcriptional complex will be tackle later in this article.

When plotting the 3’-nt of co-transcriptional processed fragments of BTL pri-miRNAs we found a clear and narrow peak coincident with the nt right before the DCL1 cleavage site (Figure 2D, 2F, S2B, and S2D; cyan arrows). Interestingly, we also detected a peak matching the last nucleotide of the miRNA-3p sequence (Figure 2D, 2F, S2B, and S2D, orange arrows). The analysis of the reads ending in this nucleotide indicates that the processed hairpin (the so-called pre-miRNA) is retained temporally in the co-transcriptional processing complex. Its lower detection agrees with our RIP validations, showing a reduced association of this region with the transcriptional complex after processing (Figure 2E). Altogether, our results indicate that plant miRNAs can be processed co-transcriptionally while nascent pri-miRNAs are still associated with the RNAPII.

### Processing of BTL pri-miRNAs initiates co-transcriptionally but ends in the nucleoplasm

Aiming to explore whether all different biogenesis modes (LTB, LTBs, BTL and BTLs) occurs co-transcriptionally, we sorted miRNAs by their processing direction (following the annotation by (Moro *et al*., 2018)) and analyzed each group individually by plaNET-seq profiling.

When we analyzed LTB pri-miRNAs, two clear peaks corresponding to the first and second DCL1 cuts were detected (Figure 3A and S3A, Cyan and purple arrows). The detection of both processing intermediates indicates that the entire miRNA biogenesis occurs co-transcriptionally when the pri-miRNA is still attached to the locus. Supporting this idea, a very similar profile was observed when LTBs pri-miRNAs were analyzed (Figure 3B, and S3B). The mapping revealed a clear pattern of peaks matching each DCL1 processing site for LTBs loci (Figure 3B, and S3C). Interestingly, an additional peak corresponding to the 3’ last nt of the miRNA-3p was visible in the global profile of both LTB and LTBs pri-miRNAs (Figure 3A, 3B, S3A and S3B, orange arrows). In LTB pri-miRNAs, this peak was only contributed by pri-miR156b, miR156e and miR408, while for LTBs, only miR159a and b contained such additional peak (Figure 3C, 3D, S3C, and S3D). Processing intermediates ending in this nucleotide (pre-miRNAs hairpin) observed before for BTL pri-miRNAs were not expected for LTB pri-miRNAs as the first processing step would prevent their existence. When we mapped the entire plaNET-seq reads over these loci, we found that all reads ending in this precise position correspond to the mature miRNA-3p (Figure 3E and F). These results show that all processing steps in LTB and LTBs pri-miRNAs occurs co-transcriptionally and that some mature miRNAs are temporally retained at their encoding loci, probably bound to a biogenesis protein such as HYL1 (Figure 3I, (Baranauske et al., 2015; Yang et al., 2010)). The observation that only some mature miRNAs derived from LTB or LTBs pri-miRNAs remain associated to their loci may represent differential affinity of each miRNA for the proteins of the biogenesis complex. It is also possible that a more efficient processing of some pri-miRNAs releases the mature miRNAs faster.

**Figure 3.**
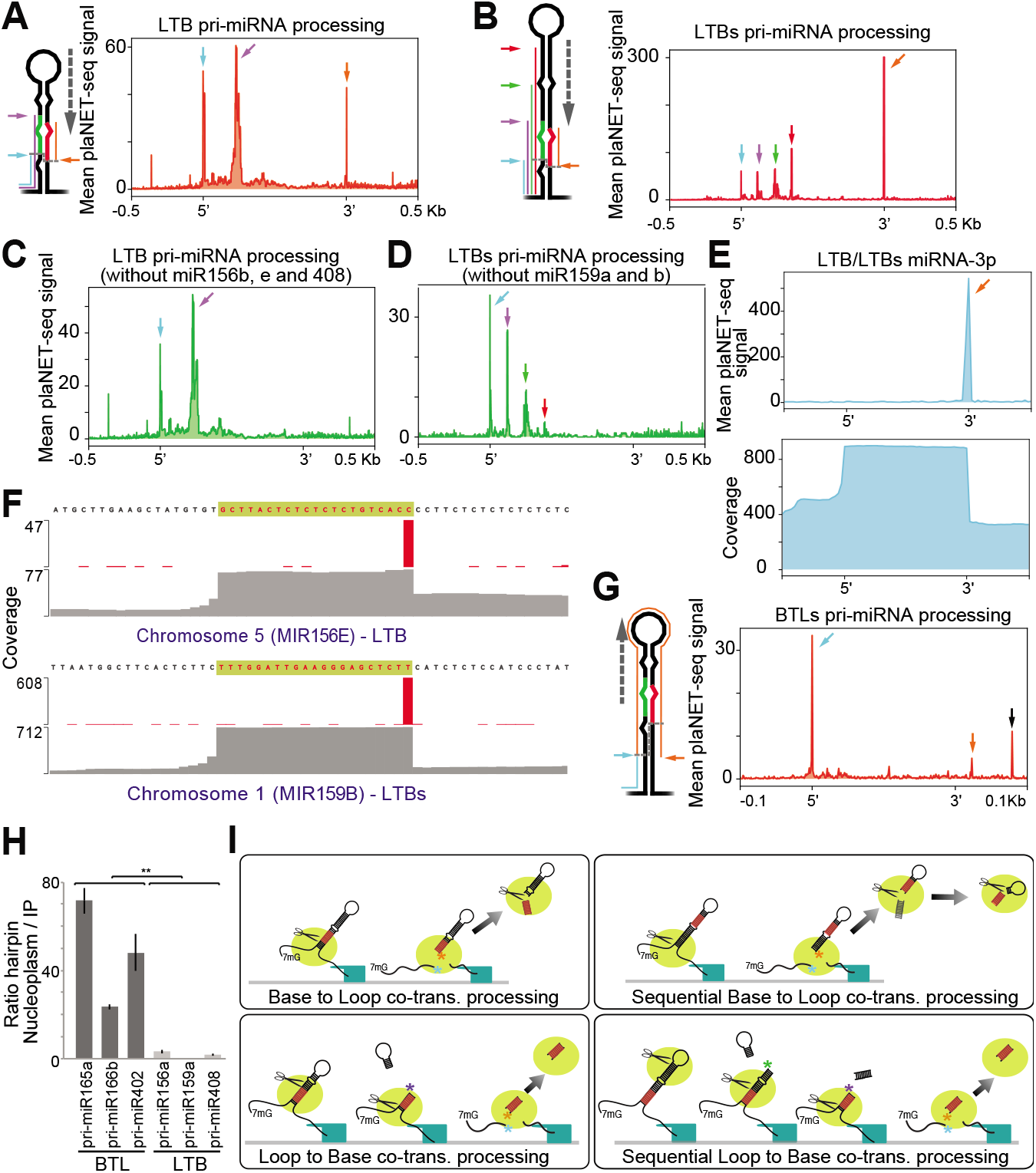
Co-transcriptional processing of BTL pri-miRNAs involves a second nucleoplasmic processing step. (A, B, C, D, and G) Metagene analysis of processing intermediated on RNAPII-nascent LTB (A and C), LTBs (B and D), and BTLs (G) pri-miRNAs as determined by scoring the 3’-end nucleotide of plaNET-seq reads. Pri-miRNAs were scaled from the beginning of miRNA-5p to the end of miRNA-3p. Colored arrows indicated the processing fragments detected for each processing type as illustrated in the schematic representation of LTB, LTBs, and BTLs pri-miRNAs. A peak corresponding to an exon donor site of *AT1G12290* is noted with a black arrow in panel (G) and Figure S3F. (E) Metagene analysis of plaNETseq LTB and LTBs scaled using the miRNA-3p sequence (top panel). Orange arrow matches the arrow in panel (B). Metagene of entire plaNETseq reads coverage over LTB and LTBs *MIRNA* loci, at the miRNA-3p encoding regions, showing the accumulation of 21-nt long reads corresponding to the miRNA-3p mature sequences (bottom panel). (F) Mean plaNET-seq signal at the *MIR156E* and *MIR159B* loci (Red bars). Coverage of plaNET-seq reads over the same loci (Grey). The sequence corresponding to the mature miRNA-3p is noted on top of each panel highlighted in yellow. (H) Relative abundance of BTL and LTB pri-miRNA hairpin region in the nucleoplasm compared to the RNAPII RIP fraction as measured by RT-qPCR in Col-0 samples. Error bars correspond to 2xSEM. P<0.01 (**), in a two-tailed unpaired T-Test, were considered statistically significant. Processed hairpins measurements in the nucleoplasm and RNAPII IP fraction were normalized to the input levels for each samples. Quantifications were then expressed as a nucleoplasm/IP ratio of the values corrected by relative amount of unprocessed pri-miRNAs quantified with primers flanking the DCL1 cleavage site. (I) Schematic summary of the co-transcriptional processing mechanisms of Arabidopsis pri-miRNAs. Color asterisks in each panel denote the plaNET-seq scored nucleotides for each processing type, and match the arrows of the same colors displayed in the previous panels and in Figure 2.

When BTL and BTLs pri-miRNAs were analyzed, we observed a peak matching the nucleotide right before the first DCL1 cleavage site but not the subsequent sites (Figure 2D, 3G, S2B, and S3E). These profiles support a scenario where only the first processing step of BTL and BTLs pri-miRNAs occurs co-transcriptionally (Figure 3I). In agreement with this hypothesis, the analysis of individual pri-miRNAs revealed the partial retention of the entire processed hairpin, the so-called pre-miRNAs (Figure 2D and 3G, orange arrows), but never the mature miRNAs as it was observed for LTB and LTBs pri-miRNAs (Figure 3A-E). These results indicate that BTL and BTLs pri-miRNAs suffer a first co-transcriptional processing step, but the resulting pre-miRNA is further processed in the nucleoplasm (Figure 3I). To confirm this hypothesis, we purified nucleoplasm and chromatin and quantified by RT-qPCR the abundance of the hairpin region relative to the unprocessed pri-miRNA in each fraction for LTB and BTL pri-miRNAs. Confirming our previous observation, we found that the hairpin regions of BTL, but not LTB, pri-miRNAs were enriched in the nucleoplasm compared to the chromatin (Figure 3H). This result supports a scenario where BTL/BTLs pri-miRNAs are processed stepwise from pri- to pre-miRNA co-transcriptionally and from pre-miRNA to miRNA duplex, post-transcriptionally. An additional peak was detected downstream the 3’-nt of the miRNA-3p (Figure 3G, black arrow) corresponding to the donor site of the first exon of an AT1G12290 splicing isoform, which also contains the miR472 encoding sequence (Figure S3F). In all analyzed cases mock-IP plaNET-seq sample (negative control) showed no peaks corresponding to processing intermediates supporting that the detected signal corresponds to processed nascent RNAPII pri-miRNAs (Figure S3G). In addition, the analyzed datasets allowed us discover that miR161.1 and miR161.2 are unique cases of dual LTB and BTL processing from the same precursor (Figure S3H and S3I). We also used the plaNETseq data to score the processing direction of previously undefined miRNAs (Moro *et al*., 2018). We defined miR157d as LTB, miR2111b as BTL with retention of the mature miRNA, and miR846 as LTBs (Figure S3J). These results indicate that it is possible to use plaNET-seq to identified processing mechanisms of pri-miRNAs in different plant species, growth conditions, or mutant backgrounds.

### Most pri-miRNAs are processed both co-transcriptionally and post-transcriptionally

The detection of co-transcriptional processing does not necessarily imply that all miRNAs, or even not all pri-miRNAs transcripts from a single locus, are processed exclusively during transcription. Full-length pri-miRNAs can be found in the cells and even move to the cytoplasm to translate into small peptides (Lauressergues et al., 2015). Thus, some pri-miRNAs, or at least a fraction of the transcripts from each locus, escape co-transcriptional processing. This is a scenario similar to splicing, where both co-transcriptional and post-transcriptional mRNA processing co-exist. To evaluate the extent of co-transcriptional processing, we re-analyzed plaNET-seq data and calculated the ratio between those reads ending at DCL1 cleavage site (co-transcriptionally processed, Figure 4A, green lines) and those expanding the site (unprocessed pri-miRNAs, Figure 4A, blue and cyan lines). Although, unprocessed pri-miRNAs associated with the chromatin will either be processed in the nucleoplasm or exit the nucleus unprocessed, we will refer to these molecules as post-transcriptionally processed pri-miRNAs. In the case of BTL or BTLs pri-miRNAs, we did this calculation using the signature peak at the 5’-end of the transcript (Figure 4A, site “a”), since the subsequent cuts are undetected and happen post-transcriptionally, as we showed before. Conversely, we used the first cleavage site toward the loop region for LTB and LTBs (Figure 4A, site “a”), as the following sites overestimate the unprocessed reads by counting processing intermediates (cyan lines). To simplify the analysis, we excluded those *MIRNA* loci without plaNET-seq mapped reads. We used two independent experiments and plotted the co-transcriptional vs post-transcriptional processing ratio for each pri-miRNA sorted by processing type (Figure 4B). In the tested conditions, and among the pri-miRNAs showing both processing components, the analysis revealed that ∼25% of the pri-miRNAs were preferentially co-transcriptionally processed, with miR166b miR165a, miR393a, miR162a, and miR160b standing out as primarily processed co-transcriptionally (Figure 4B-D). Approximately 8% of analyzed miRNAs showed an equal preference for co-transcriptional and post-transcriptional miRNA processing (Figure 4B-D). While ∼67% of pri-miRNAs appeared to be processed more post-transcriptionally (Figure 4B-D). From those, it is worth noting that ∼31 % (21% among all) have a ratio > to 0.5, indicating that those miRNAs are processed co-transcriptionally more than a quarter of the time (Figure 4B-D). Several pri-miRNAs, including those encoding miR862, miR399b, miR165b, and miR394b, stand out among those with a nearly undetectable signal of co-transcriptional processing (Figure 4B). These results suggest that pri-miRNAs co-transcriptional processing is likely a dynamic and potentially regulated process, not identical for each locus.

**Figure 4.**
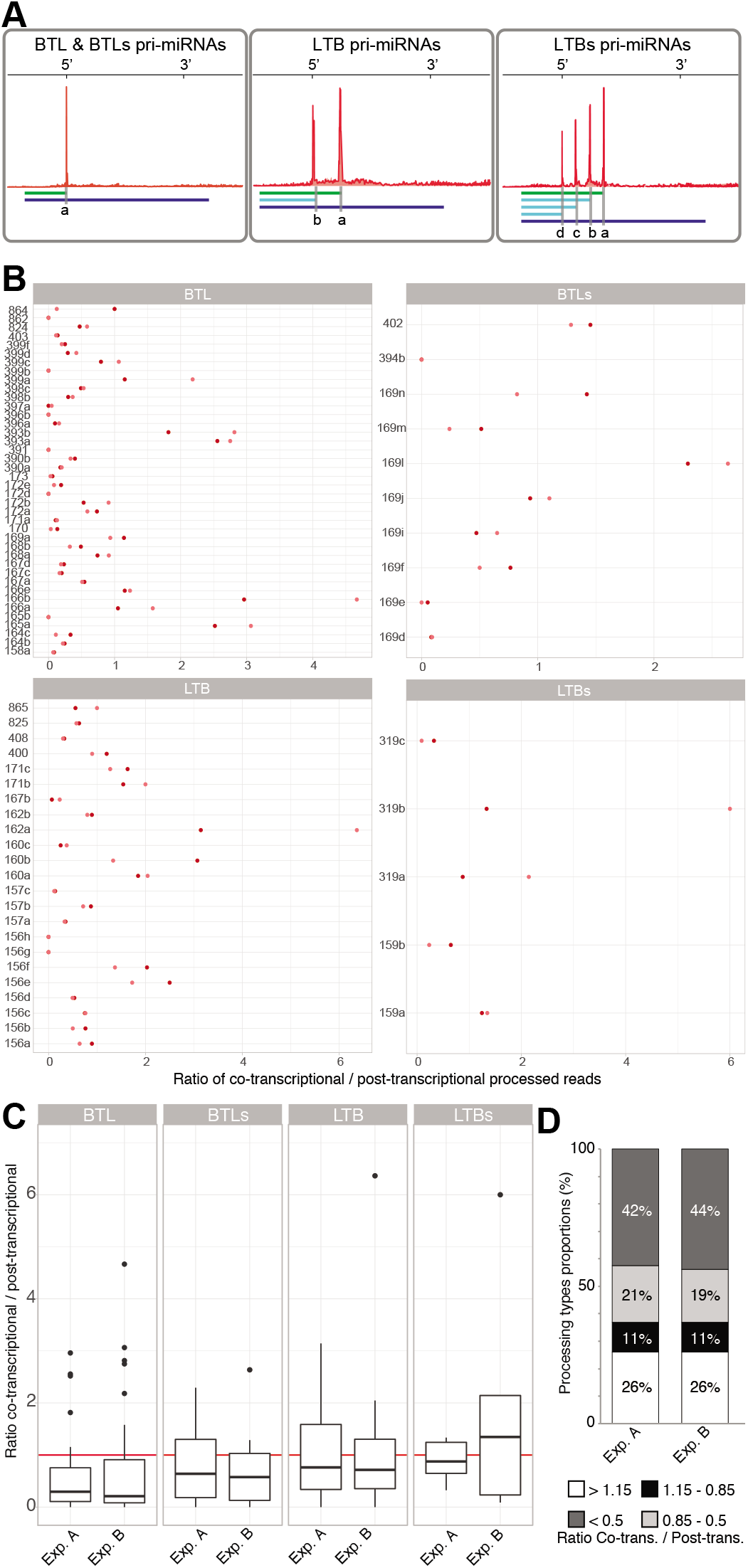
Co-transcriptionally and post-transcriptionally processing co-exist for most pri-miRNAs. (A) Schematic representation of the positions used to quantify co-transcriptional vs post-transcriptional processing ratios. In all cases total number of reads ending in the nucleotide marked as (a) (green lines) were expressed relative to reads expanding this site (blue lines). (B) Co-transcriptional processing ratios corresponding to all analyzed miRNAs in two independent plaNET-seq experiments and split by processing mechanism. (C) Box-plot representation of the co-transcriptional processing ratios. Red line mark ratio =1 where co- and post-transcriptinal processing are equally frequent. (D) Fraction of pri-miRNAs preferentially processed co-transcriptionally, post-transcriptionally or with equal preference. Two independent experiments in control conditions and the addition of both are displayed.

### Co-transcriptional miRNA processing efficiency and frequency fluctuate depending on the conditions

We next wondered whether the transcriptional activity of the RNAPII or even the environment could affect the balance between co-transcriptional and post-transcriptional processing of pri-miRNAs. We first repeated our previous analyses using plaNET-seq data obtained from plants incubated for 12 hours at 4 °C. The results indicated that reducing the plant growing temperature impact the ratio between co-transcriptional vs post-transcriptional processing (Figure 5A), especially for some individual miRNAs (Figure S4A, S4B). Among all tested pri-miRNAs, a comparison of the co-/post-transcriptional processing ratios between samples (with a threshold of +/− 50% difference) underlines a considerable number of pri-miRNAs affected by this condition either in a negative or positive way (Figure S4B).

**Figure 5.**
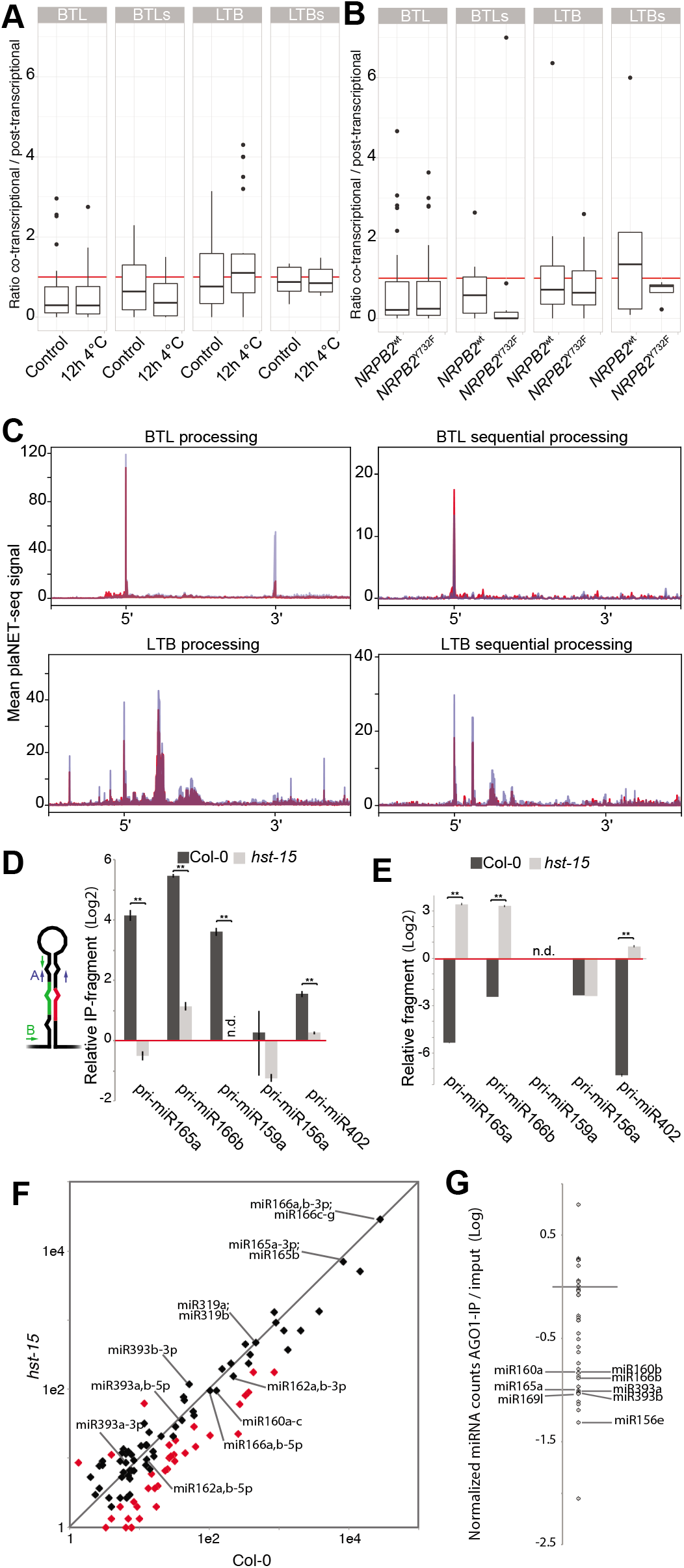
Co-transcriptional miRNA processing ratios are variable in different conditions. (A and B) Box-plot representations of the co-transcriptional processing ratios in seedlings transfer to 4 °C for 12 hours or kept at control conditions (A) or *nrbp2* mutant plants expressing a WT version of the proteins or a mutation (Y732F) that confers enhanced processivity to the RNAPII (C). Red line mark ratio =1 where co- and post-transcriptinal processing are equally frequent. (E) Superposition of metagene analysis of plaNET-seq pri-miRNA processing intermediated in control (red) or NRPB2^Y732F^ transgenic plants (Blue). Pri-miRNAs were scaled from the beginning of miRNA-5p to the end of miRNA-3p. (F, and G) Pri-miRNA co-transcriptional processing in Col-0 and *hst-15* mutant plants as measured by RT-qPCR in RIP (F) or chromatin depleted nucleoplasm samples (G). Co-transcriptional processing was measured as the relative abundance of the hairpin region (Primers A) over the amount of unprocessed pri-miRNAs quantified with primers flanking the DCL1 cleavage site (Primers B). Processing intermediates were normalized by the input and expressed relative to the IgG IP samples (red line). Error bars corresponds to SEM. P<0.01 (**), in a two-tailed unpaired T-Test, were considered statistically significant. (H) Scatter plot comparing the counts per million +1 (log scale) of mature miRNAs between wild type (Col-0) and *hst-15* mutants. Differentially expressed miRNAs are shown in red. MiRNAs with the largest ratio of co-transcriptinal processing are noted individually. (I) Dot-plot of AGO1-loaded miRNAs. MiRNAs in the AGO1-IP fraction as well as in the input samples were first expressed as a fraction of the total count of miRNAs in the respective sample. AGO1 loading preference for each miRNA is then expressed as the ratio of the frequency of a miRNA in the IP vs the input sample. MiRNAs with the largest ratio of co-transcriptinal processing are noted individually.

The ratio of exon/intron inclusion, one of the best-studied co-transcriptional RNA maturation processes, is well known to depend on RNAPII elongation speed (Dujardin et al., 2014; Giono and Kornblihtt, 2020; Godoy Herz and Kornblihtt, 2019). Thus, we wondered whether the pri-miRNAs co-transcriptional/post-transcriptional ratio is also affected by the RNAPII activity. To test this possibility, we analyzed plaNET-seq data obtained from *nrpb2-1* plants complemented with a wild type *NRPB2:FLAG* construct (*NRPB2*^wt^) or with a version of this protein containing a Y732F mutation (*NRPB2*^Y732F^) which accelerates RNAPII transcription *in vivo* (Leng *et al*., 2020). Using a difference ratio threshold of 0.5, we found that, with a few exceptions, an increase in RNAPII speed produces a reduction in co-transcriptional processing. However, this is evident for a relatively small fraction (∼22%) of all analyzed miRNAs (Figure 5B, S4C, and S4D). Given that at least the entire stem-loop region of a pri-miRNA needs to be transcribed before processing starts, our result may imply that a quick transcription releases the mature pri-miRNA transcript before it is co-transcriptional processed.

Next, we analyzed the co-transcriptional processing profile in NRPB2^Y732F^ plants, asking whether RNAPII speed may affect co-transcriptional processing efficiency rather than frequency. Overall, we did not find any noticeable effect over pri-miRNAs processed BTL beyond an increment in the retention of the hairpin (Figure 5C). Interestingly, we found an apparent enhanced processing efficiency in LTB and LTBs pri-miRNAs (Figure 5C). In particular, it was interesting to observe that such increase in the processing efficiency was progressive from the initial DCL1 cut and become more evident in the subsequent cuts. This intriguing result may represent partial, alternative, or misfolded pri-miRNAs generated depending on the polymerization speed. Although, the number of LTBs pri-miRNAs is small and we cannot discard this as a coincidence.

Overall, these results suggest that the balance between co-transcriptional and post-transcriptional processing of each pri-miRNA is dynamic and responds to specific conditions. This opens the question of whether a miRNA may have different functions depending on when/where it is processed. Recently, we have reported that HST associates with *MIRNA* loci through the interaction with the mediator complex, recruiting DCL1 to the *MIRNA* loci (Cambiagno *et al*., 2021). Thus, we reasoned that HST mutants might have an imbalance of the processing ratio that can help us study the dynamics of co-/post-transcriptional processing. We measured the amount of processed and un-processed pri-miRNAs in the nucleoplasm and chromatin of Col-0 and *hst-15* mutant plants to test this hypothesis. Confirming *hst-15* as a model of impaired co-transcriptional processing, we found a reduction in the ratio of processed/non-processed nascent pri-miRNAs in the IP fraction in the mutants compared to WT plants (Figure 5D). We observed the opposite pattern in the nucleoplasm fraction, suggesting that the balance between co-transcriptional and post-transcriptional processing swoop in this mutant, but the overall miRNA production is compensated (Figure 5E). Coincidentally, small RNA sequencing analysis of *hst-15* mutants revealed that most miRNAs, and particularly those with high co-/post-transcriptional ratio, are not altered in the mutants as previously reported (Brioudes et al., 2021), supporting a change in processing type rather than an overall effect on miRNA biogenesis (Figure 5F). Recently, it was shown that HST mutants display a compromised non-cell-autonomous miRNA function cause by an impaired movement of mature miRNAs (Brioudes *et al*., 2021). Notoriously, most miRNAs reported to act non-cell-autonomously, such as miR160, miR165, and miR166 (Brioudes *et al*., 2021; Brosnan et al., 2019; Fan et al., 2021), ranked among the highest co-transcriptionally processed miRNAs in our data (Figure 4B). In agreement with the reports suggesting that AGO1-unloaded miRNAs are likely the mobile component (Brioudes *et al*., 2021; Devers et al., 2020; Fan *et al*., 2021), highly co-transcriptionally processed miRNAs appeared among the less efficiently loaded in AGO1 in RIP-seq experiments (Figure 5G). This suggests that co-transcriptionally processed miRNA may undergo a different fate after processing. This idea goes in line with the reduced co-transcriptional miRNA processing observed in *hst-15* (Figure 5D) and the lack on miRNA movement, without a change in the mature miRNA steady levels, previously reported for this mutant (Brioudes *et al*., 2021).

### R-Loops formation between pri-miRNAs and the encoding loci promote co-transcriptional miRNA processing

One of the most intriguing results so far was the systematic detection of the processed 5’ region of the pri-miRNAs in the NRPB2-IP fraction. Similar profiles were previously reported, for example, during splicing due to the stabilization of the processed mRNA fragments by the spliceosome complex (Nojima *et al*., 2015). Different from splicing, where each intron can be removed while transcription proceeds, co-transcriptional pri-miRNA processing required the transcription of at least the entire stem-loop region before the processing complex can recognize the features necessary for DCL1-mediated miRNA biogenesis. Thus, it is possible that the observed retention of the 5’ ssRNA arm in the IP samples represent pri-miRNA transcripts either stabilized or anchored to the loci to become co-transcriptionally processed. The observation in our microscopy assays that spliced pri-miRNAs are still associated to their transcription sites (Figure 1) also support a scenario were pri-miRNAs are temporally anchored to the encoding loci. The apparent effect of transcriptional speed on the co-processing efficiency made us wonder, as well, whether transcription influences such anchoring. A slow transcription may cause, for example, the formation of R-Loops between an RNA and its encoding locus, which in turn could lock the transcript in the locus (Zatreanu *et al*., 2019). Within such a scenario, an R-loop that is formed between a *MIRNA* encoding locus and the nascent transcript may stabilize and anchor the pri-miRNA to allow co-transcriptional processing once the transcription and folding of the dsRNA hairpin structure end. We explored this possibility by analyzing DRIP-sequencing (DRIP-seq) data (Xu et al., 2020; Xu *et al*., 2017), searching for the presence and pattern of R-Loop formation over *MIRNA* loci. To do this analysis, we first scaled all pri-miRNAs using either a fixed window from the transcription start site (TSS) to the most up-stream DCL1 cleavage site (Figure 6A, green line) or from this site to the end of miRNA-3p (Figure 6A, orange line). To establish such windows, we first defined each pri-miRNA TSS as described in Materials and Methods. Then, we scored the R-Loop signature for each pri-miRNA, and plotted the metagene profile of R-loops over *MIRNA* loci (Figure 6B). We found frequent R-loops over the analyzed loci in the DRIP-seq dataset, an observation confirmed by DRIP-qPCR on some specific loci (Figure 6B, and 6C). However, it was clear that the pattern of R-loop formation was variable among tested pri-miRNAs. Thus, we sorted all pri-miRNAs into categories depending on their R-Loop profile (Figure 6D). We observed five types of *MIRNA* genes: those not showing R-loops at all (Figure 6D (α)), those with R-loops restricted to the 5’-ssRNA arms of the pri-miRNAs (Figure 6D (β)), those with an R-loop like β but with an additional signature toward the end of the loci (Figure 6D (γ)), those where the R-Loops extend uniformly over the entire loci (Figure 6D (δ)) and finally miRNAs with colliding R-loops, probably a reflection of bi-directional transcription (Figure 6D (ε)). Strikingly, when we asked whether the presence of an R-Loop impacts co-transcriptional processing, measured as the ratio co-/post-transcriptional processing, we found that miRNAs with R-Loops in the initial single stranded region of the transcripts (β and γsignatures) are preferentially processed co-transcriptionally (Figure 6E). Conversely, pri-miRNAs not displaying R-loops over the loci or the characteristic R-Loop near the TSS (α or δ,ε respectively) showed poor signs of co-transcriptional processing (Figure 6E).

**Figure 6.**
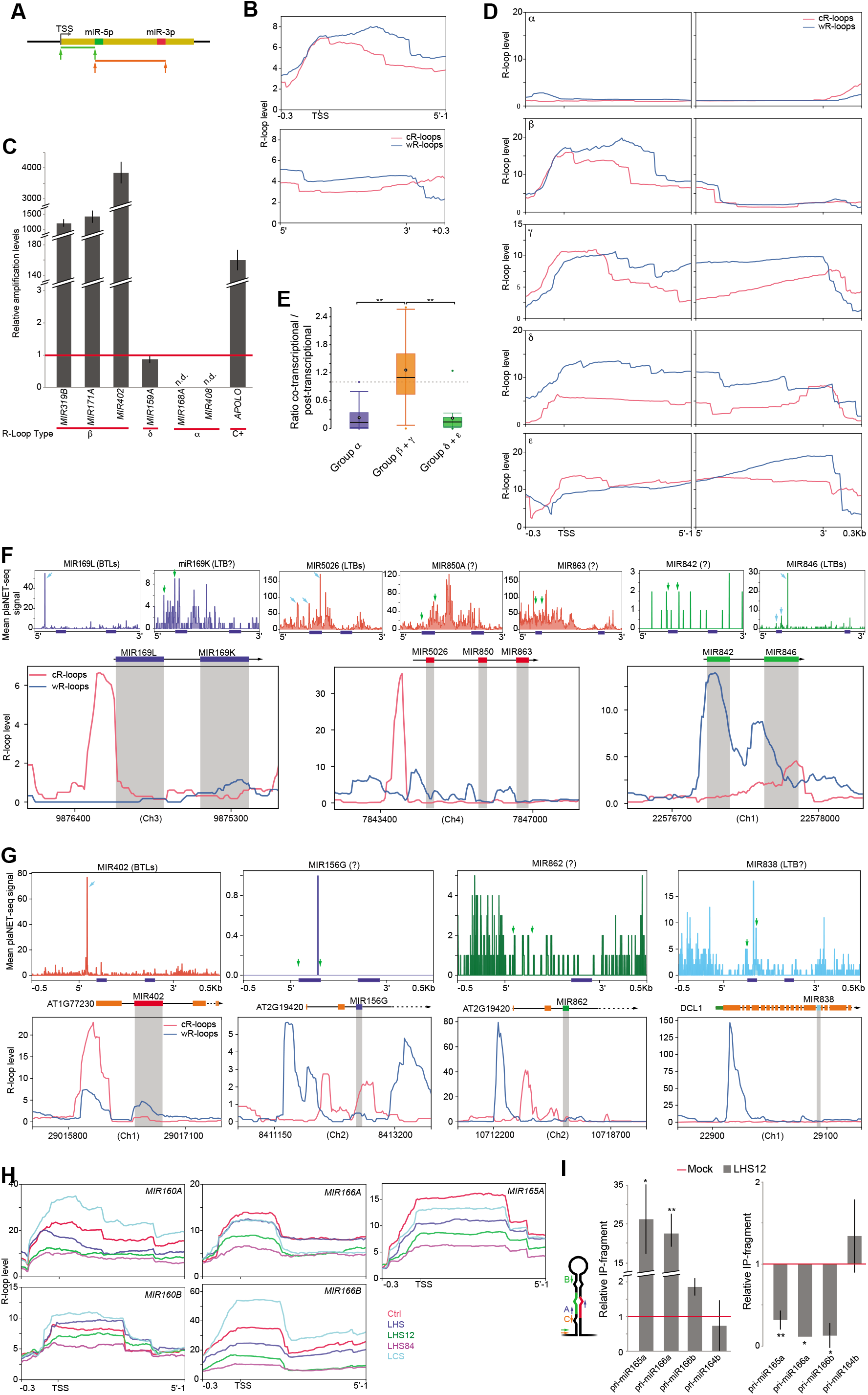
R-Loops at the 5’ end of miRNA loci promotes co-transcriptional processing. (A) Schematic representation of the region scaled for metagene analysis of R-Loop formation over *MIRNA* loci. *MIRNA* loci were scaled either from the transcription start site (TSS) to the nucleotide before miRNA-5p (green) or from miRNA-5p to the end of miRNA-3p (red). Metagene analysis was then plotted in (B and D) adding 300 bp upstream or downstream the windows respectively. (B) Metagene analysis of R-loop formation over all MIRNA loci either in the Watson (blue) or Crick (red) strands. (C) Quantification of R-loops near the TSS of *MIRNA*s belonging to different categories of R-loop pattern (defined in (D)) as measures by DRIP-qPCR assays. Values of DRIP samples are normalized to the input and expressed as relative to the DRIP signal in samples IP with a α-IgG antibody (red line). Failure to detect R-Loops are notes as non-detected (n.d.). Error bars correspond to 2xSEM. (D) Metagene analysis of R-loop formation over *MIRNA* loci sorted by R-loop distribution. α-*MIRNA* loci without R-loops; β- *MIRNA* loci with R-loops over the ssRNA 5’ end of the pri-miRNAs; γ- *MIRNA* loci with bipartite R-loop signals at the beginning and end of the pri-miRNAs; δ- *MIRNA* loci with R-loops over the entire loci; ε- MIRNA loci with colliding R-loops in the Watson (blue) and Crick (red) strands. (E) Boxplot of co-transcriptional processing ratios of pri-miRNAs with R-loop profiles of categories α, β + γ, or δ + ε. Equal processing type frequency (=1) is marked with a dashed line. Error bars shows the maximum and minimum values. P<0.01 (**), in a two-tailed unpaired T-Test, were considered statistically significant. (F, and G) R-loop profile and plaNET-seq signals on polycistronic miRNA clusters (F) and mirtron loci (G). Cyan arrows in the plaNET-seq plots indicate accurate detected processing site. Green arrows mark the positions where peaks would be expected if the corresponding pri-miRNA are co-transcriptionally processed. The position of miRNA-5p and −3p are marked with blue boxed under the plaNET-seq plots. MiRNA precursor sequences within the containing locus are noted in gray within the R-loop profiles. (H) R-loop profile over individual miRNA loci as detected in samples extracted from control plants (Red line), plants incubated 30 h at 17 °C (LCS) or at 37 °C (LHS), plants incubated 30 h at 37 °C and then returned to 23 °C for 12 (LHS12) or 84 (LHS84) hours. (I) Left panel: Co-transcriptional processing ratio in the H3 RIP fraction as measured as the accumulation of the hairpin region (Primers A) normalized to the unprocessed pri-miRNA (Primers B) by RT-qPCR. Right panel: Retention levels of processed 5’-end arms of pri-miRNAs (primers C) in the IP sample normalized by the input sample as measured by RT-qPCR. In both panels the quantification in the plants incubated for 30h at 37 °C + 12h at 12°C (LHS12, grey bars) are expressed relative to the control samples (red line). Error bars corresponds to SEM. P<0.01 (**) and P<0.05 (*), in a two-tailed unpaired T-Test, were considered statistically significant.

Remarkably, when we analyzed polycistronic miRNA duets encoded by a single transcript, we observed that the miRNA hairpin located adjacent to an R-loop was processed co-transcriptionally more efficiently or even the only one showing signs of co-transcriptional processing (Figure 6F). This goes in line with a recent report indicating that polycistronic 5’ hairpins are processed more efficiently than 3’ hairpins (Lunardon et al., 2021). It also reinforces the hypothesis that R-loops between nascent pri-miRNAs and its encoding loci anchor and stabilize the molecule allowing the co-transcriptional processing. Interestingly, in the case of MIR842/MIR846 duet, the second miRNA, but not the first, shows signs of co-transcriptional processing (Figure 6F). This is coincident with an extended R-loop encompassing the entire region encoding MIR842, likely impairing its processing as observed for class δ *MIRNAs*, but stabilizing the second hairpin to allow miR846 co-transcriptional processing as β *MIRNAs*.

When we explored some miRNAs encoded within protein-coding genes, we found that miR402, which is processed form a short transcript (Knop et al., 2017) and encoded in the first intron of the hosting gene near the TSS and associated with an R-Loop, is efficiently processed in a co-transcriptional way (Figure 6G). Conversely, miRNAs encoded away from the TSS and the associated R-loops, such as miR156g, miR862 or miR838, do not show co-transcriptional processing signs or only an inefficient processing (Figure 6G). This observation reinforces the idea that R-loops in the proximity and upstream of the hairpin promotes co-transcriptional processing of pri-miRNAs.

Interestingly, parallel studies using similar, but not identical, samples (Xu *et al*., 2020; Xu *et al*., 2017) showed different R-loop patterns in some miRNA loci (Figure S5). This suggest that R-loop formation may change depending on the plant developmental stage or growth condition and potentially regulate co-transcriptional miRNA processing. We mined DRIP-seq data (Xu *et al*., 2020) searching for growth conditions with altered R-loop formation patterns to explore further whether R-loops initiate and regulate co-transcriptional processing. We focused only on those *MIRNA* loci with significant changes in the processing ratio (Figure 5). An alteration in the R-loop patterns may represent a regulatory mechanism that translates into changes in the processing mechanisms and potentially on the mature miRNA fate. Among the tested condition, we found that prolonged heat stress (30 hours at 37 °C), either control or after a period of recovery at 23 °C, produced consistent changes in the R-Loop patterns over MIR160a, MIR160b, MIR165a, MIR166a, and MIR166b among others (Figure 6H). In agreement with the role of R-loop in promoting co-transcriptional processing, we observed that the same temperature treatment also translated in a reduction of the co-transcriptional processing ratio measured as an increment in unprocessed pri-miRNAs in RIP-qPCR samples (Figure 6I, left panel). This change in co-transcriptional processing was not observed for pri-miR164b, which does not present an R-loop over its locus and showed undetectable signs of co-transcriptional processing (Figure 6I, left panel). In addition, we also found that the processed ssRNA arm of pri-miR165a, 166a, and 166b, but not 164b used as a negative control, were less retained in the RIP fraction of heat stressed plants coincident with the reduction in R-loops (Figure 6I, right panel). These results position the co-transcriptionally formed R-loops as a critical element to anchor and trigger co-transcriptional processing of pri-miRNAs in plants. This also suggested that the regulation of R-loops resolution directly impact how miRNAs are produced. Even when it is likely that the miRNA biogenesis complex also helps to stabilize the nascent pri-miRNAs in the transcriptional complex, the formation of R-loops appeared as a critical event toward achieving co-transcriptional miRNA processing.

## Discussion

The coupling of transcription and RNA processing is a common feature in most organisms. Nascent RNAs undergo several processing steps co-transcriptionally, including 5’-capping, splicing, polyadenylation, as well as chemical modifications such as m6A (Bentley, 2014; Lee and Tarn, 2013; Yang *et al*., 2021). In animals, the processing of pri-miRNAs is not an exception and is accepted to occur co-transcriptionally (Morlando *et al*., 2008; Nojima *et al*., 2015; Pawlicki and Steitz, 2008). However, it was unclear whether this process also occurs co-transcriptionally in plants despite many miRNA-biogenesis factors associate with *MIRNA* loci. The long and structurally variable plant pri-miRNAs posed a challenge for such an event to happens. On top, the recent demonstration of pri-miRNA processing in SERRATE-containing liquid droplets, likely D-Bodies, and the absence of this protein in most *MIRNA* loci challenged the idea of co-transcriptional processing in plants (Speth et al., 2018; Xie et al., 2021). In this study we confirmed that pri-miRNAs are processed co-transcriptionally in plants and showed that this process co-exists with a post-transcriptional counterpart. This implies that two alternative pathways, perhaps involving a different set of proteins, co-exist and may produce miRNAs with alternative functions. We have shown that HST is required for DCL1 association to *MIRNA* loci and pri-miRNA co-transcriptional processing (Cambiagno *et al*., 2021), Figure 5F, and G). Recently, it was also demonstrated that HST is required for the non-cell-autonomous function of miRNAs (Brioudes *et al*., 2021). However, it is unclear the mechanism involved in such an HST-depended miRNA-movement. Based on the results presented here, it is tempting to speculate that co-transcriptionally processed miRNAs, but not their post-transcriptionally processed siblings, constitute the mobile pool of miRNAs. It is possible, for example, that AGO1 preferentially loads post-transcriptionally processed miRNAs sealing their fate as non-mobile molecules. In this scenario, the interaction of AGO1 with nucleoplasmic exclusive miRNA partners, such as CARP9 or TRANSPORTIN1 (Cui et al., 2016; Tomassi et al., 2020), but not with the chromatin-associated complex, may sort which miRNAs are loaded. On the other hand, co-transcriptionally processed miRNAs may scape nuclear AGO1-loading by missing partner proteins, shuttling to the cytoplasm either free or associated with chaperon proteins, such as HYL1, and then become mobile. However, it was reported that AGO1 could also interact with the chromatin and even RNAPII (Huang et al., 2013; Liu et al., 2018) suggesting that the outcome of co-transcriptional processed miRNA may be unique for each loci. It is worth noting that miRNAs know to act non-cell-autonomously, such as miR160, miR165, and miR166, rank on top of the most co-transcriptionally processed miRNAs in our analysis (Figure 4). Interestingly, our study showed that the preference for co- or post-transcriptional processing swoop for mobile miRNAs in *hst* mutants which translate in nearly unaffected mature miRNA levels the mutants as previously reported (Brioudes *et al*., 2021) (Figure 5F-H).

Two proposed models explain co-transcriptional modifications of RNAs in animals and yeast. A first model, known as the recruitment model, relies on the transcriptional machinery to recruit RNA processing factors to trigger the events (Bentley, 2014). This model appears relevant during the coupling of transcription and pri-miRNA processing in plants as many miRNA biogenesis factors rely on the interaction with the RNAPII transcriptional complex to associate to *MIRNA* loci. In a second model, the kinetic model (de la Mata et al., 2010), the relative rates of transcription elongation directly impact RNA processing. This is the case of splicing or poly(A) sites that are recognized or skipped depending on the RNAPII speed producing alternative transcript isoforms. In this process, a slow elongation rate provides RNA processing factors with more time to recognize processing sites, to assemble the complexes, and to produce the modifications (Dujardin *et al*., 2014). This model could also be particularly relevant for plant pri-miRNAs co-transcriptional processing, as the long transcript requires time, and a large percentage of transcription completion to fold correctly. It was recently shown that the transcription elongation rate impacts nascent RNA folding (Saldi et al., 2018; Scharfen and Neugebauer, 2021). Thus, a change in the elongation rate may affect the structure of pri-miRNA folding and how it is processed, as previously shown (Wang et al., 2018). Interestingly, N^6^-adenosine m6A methylation, a co-transcriptional RNA modification (Knuckles et al., 2017), was show to impact RNAPII pausing (Akhtar et al., 2021) and pri-miRNA processing (Bhat *et al*., 2020) providing another potential link between co-transcriptional events and miRNA biogenesis. In this sense, our data indicated that elongation speed affect co-transcriptional processing, perhaps by controlling R-loop formation (Figure 5). However, such a potential effect needs to be studied case by case rather than in a metagene analysis as each pri-miRNA structure is unique and affected differently.

RNAPII speed (elongation rate) is regulated in response to intra- and extra-cellular stimuli changing transcriptome composition in turn (Muniz et al., 2021). Interestingly, R-loops are formed during transcription and impact RNAPII elongation (Aguilera and Garcia-Muse, 2012). In turn, slow elongation by RNAPII also increases co-transcriptional R-Loop formation (Zatreanu *et al*., 2019). This implies that the stimuli the cells receive regulate R-loop formation. Here, we showed that the hybridization of the nascent pri-miRNA with their encoding loci promotes co-transcriptional processing by anchoring the 5’ ssRNA arm of the pri-miRNA to the loci by R-loop formation. R-loops have previously been linked to many biological processes by directly affecting transcription and genome stability. In our model, it is likely that R-loops adjacent to the TSSs of *MIRNA* loci stabilize the nascent pri-miRNA, and provide the time required to transcribe and fold the stem-loop region, and to assemble the miRNA-biogenesis complex co-transcriptionally. To the best of our knowledge, this is the first time such a positive regulatory function is attributed to R-loops as a feature improving the RNA own processing. Nevertheless, it can be argue that R-loop formation could potentially negatively impact *MIRNA* transcription as RNAPII would collide with the hybrid (Aguilera and Garcia-Muse, 2012). However, it has been shown that R-loops are dynamic structures with a rapid turned over of a half-life of 10 min (Chedin, 2016; Sanz et al., 2016). Thus R-loops are continuously formed and resolved, allowing only temporal retention of nascent transcripts, probably only sufficient to process the pri-miRNA co-transcriptionally in this case.

In summary, our study provides evidence of the existence of a co-transcriptional processing pathway in plants, a mechanism that co-exists with a canonical nucleoplasmic process. We also provide new insights into mechanisms of pri-miRNA processing that depends on processing direction. We found that the co-transcriptional processing of BTL miRNA resembles the animal pathways with more defined pri-miRNA>pre-miRNA>miRNA steps. Conversely, LTB pri-miRNAs follows a more fluid path with continuous processing steps that blurs the canonical stages. The discovery that R-loops between the nascent pri-miRNAs and the encoding loci promotes coupling between transcription and processing provides an novel regulatory scenario that can re-define the function of a mature miRNA. Identifying the proteins, likely RNA-helicases, which help resolving these R-loops is imperative to manipulate the ratios of co-transcriptional processing and study the potential function of the produced miRNAs.

## Material and methods

### Plant material and growth condition

*Arabidopsis thaliana* ecotype Columbia (Col-0), transgenic and mutant plants were grown at 23° C on plates containing 2.2 g/L of Murashige-Skoog (MS) medium (pH 5.7) and 0.6% agar in long-day photoperiod (LD, 16 hours of light/8 hours of dark). Seeds were disinfected with 10 % v/v bleach and 0.1 % SDS and stratified in 0.1 % agar for 3 days at 4° C before sowing. Arabidopsis thaliana seed ecotype Columbia (Col-0), *hst-15* (SALK_079290), *hyl1-2* (SALK_064863), *ATHB1-HIS* (Miguel *et al*., 2020), *NRPB2-FLAG* (Onodera et al., 2008), and *NRPB2^Y732F^-FLAG* (Leng *et al*., 2020), were used in this study. For isolation chromatin/nucleoplasm RNA experiments the plants were grown for 20 days under before collecting the samples. For long heat stress (LHS) treatments the seeds were grown in ½ - strength MS medium complemented with 0.6% agar for 12 days at 23 °C, then the seedlings were transferred to 37°C for 30 hrs. before returning then to 23 °C for 12 h. For FISH experiments the plants were grown on Jiffypots® (Jiffy-7 42 mm; Jiffy Products International AS, Norway) and stratified for 2 days in dark at 4 °C. The immunolabeling of BrU and α-amanitin treatment experiments were performed on isolated nuclei of 2-week-old seedlings grown on ½ - strength MS medium complemented with 0.8 % agar. Before isolation, the plants were fixed in 4 % paraformaldehyde in phosphate-buffered saline (PBS), pH 7.2.

### Isolation chromatin/RNAPII-bound RNA and nucleoplasm RNA

Between 3-4 grs of plant material was frozen in nitrogen liquid and grinded in a mortar. The powder was resuspended in 30 mL of Extraction Buffer 1 (10mM Tris-HCl pH8; 0.4M sucrose; 10 mM MgCl_2_; 5 mM BME; 0.2 mM PMSF; RNasin PROMEGA) and was filter thought a Nylon membrane of 150 μm and centrifuged at 2000 g for 20 min at 4 °C. The pellet was washed twice time with Extraction Buffer 2 (10mM Tris-HCl pH8; 0.25 M sucrose; 10 mM MgCl_2_; 5 mM BME; 1 % TRITON X100; 100 μM PMSF) and centrifuged at 2000 g for 10 min at 4 °C. Then we added 500μl of Extraction Buffer 3 (10mM Tris-HCl pH8; 1.7 M sucrose; 2 mM MgCl_2_; 5 mM BME; 0.15 % TRITON X100) to the pellet. This solution was gently placed on a 1500 μl column of Extraction Buffer 3 and centrifuged at 13000 g for 5 min at 4 °C. The pellet obtained was resuspended in 500μl Lysis Buffer (0.3 M NaCl; 20 mM Tris-HCl pH7.5; 5 mM MgCl_2_; 5 mM DTT; proteases inhibitor tablet) and incubated at 4°C for 2 h in a rotator. We took 10% of the sample and saved it as INPUT, 45% for IgG-IP (AS09 605, Agrisera) negative control, and 45% used for the RIP experiment. For this, 30 μl of SureBeads (Protein A Magnetics Beads, BioRad) and 1/1000 of Histone 3 (H3 AS10 710, Agrisera) or RNAPII (AS11 1804, Agrisera) antibody were added to the sample and incubated in rotation at 4°C overnight. The IP fraction was saved as RIP-sample while the supernatant as nucleoplasm. The RIP fraction was washed with Washing Buffer (0.3 M NaCl; 20 mM Tris-HCl pH7.5; 5 mM MgCl_2_; 5 mM DTT; protease inhibitor tablet; RNasin PROMEGA) three times. Treatment with Proteinase K was carried out in 500 μl of PK Buffer (100 mM Tris-HCl pH8; 50 mM NaCl; 10 mM EDTA; Proteinase K 4 mg/ml; RNasin PROMEGA) 2 h at 55 °C and 15 min at 95 °C. We added 2U of DNase I (Thermo Fisher) to the sample and incubated them for 30 min at 37 °C. RNA extraction was then performed with 1 ml of TRIZOL and 200 μl of chloroform. Precipitation was done with 1 μl glycogen, acetate of sodium 3 M pH2.5, and isopropanol at −20°C overnight. The reverse transcription was performed with EasyScript Reverse Transcriptase (M-MLV, RNase H-, TransGen Biotech) and dN6 according to the manufacturer recommendations. Quantitative RT-qPCRs, were performed using three independent biological replicates. U6 was used as a housekeeping loading control. Averages from biological replicates and SEM were calculated from 2^-ΔΔCt^ values, and the error displayed as two times SEM. Each replicate was treated as independent samples for statistical analysis. Statistical differences between samples were determined by an unpaired, two-tailed, t-test analysis. See supplementary table S1 for oligonucleotide primers.

### DNA-RNA Immunoprecipitation (DRIP)

This experiment was performed with 3 grams of plant material previously frozen in nitrogen liquid. After grinded of samples, we performed the nucleus purification as described above. The chromatin pellet was resuspended in 300 μl Nuclei Lysis Buffer (50 mM Tris-HCl pH8; 0.1 % SDS; 10 mM EDTA). We added 300μl of Proteinase K Buffer 2X (200 mM Tris-HCl pH7.5; 100 mM NaCl; 20 mM EDTA; RNasin PROMEGA 20 U/μl; 0.04 mg/ml Proteinase K) and incubated for 1 h at 55 °C. We then added 1 volume of Phenol-Chloroform-Isoamyl acid (25:24:1) solution, mixed and centrifuged for 15 min at 12000 rpm at 4°C. We took the upper phase and transferred it to a new tube. We added 1 volume of chloroform, mixed and centrifuged for 15 min at 12000 rpm, 4°C. We sonicated the upper phase in Refrigerated PicoRuptor for 4 cycles, 30 seconds ON – 30 seconds OFF. In this step, the sample was divided into three fractions: 10% of the sample was used as an INPUT, 45% for RNase H treatment as negative control, and 45% for DNA-RNA immunoprecipitation with S9.6 antibody (MABE1095 Millipore-SIGMA) and Dynabeads Protein G overnight at 4 °C. The next day, the beads were washed three times with ChIP Dilution Buffer (1.1% TRITON X100; 1.2 mM EDTA; 16.7 mM Tris-HCl pH8; 167 mM NaCl) 5 min in rotation at 4°C. After the washes we resuspended the beads in 500 μl of Proteinase K Buffer 1X and incubated at 55-65°C for 1hr and at 95°C for 15min. The DNA-RNA purification was performed with Phenol-Chloroform-Isoamyl. After washing with chloroform, we added 1 μl of glycogen, 10% volume of NaAc pH5.2, and two-volume of absolute ethanol and incubated it overnight at −20°C. DNA-RNA was recovered by centrifugation for 30 min at 12000 rpm, 4°C. The pellet was washed with 300 μl of ethanol 70%. The dry pellet was resuspended with 30 μl of water supplemented with 0.5 μl of RNAse A (EN0531 Thermo-Fisher) and the resulting DNA used for qPCR analysis as described before.

### RNA analysis

The rapid amplification of 5′ cDNA ends (5′ RACE) method to detect processing intermediates was carried out from RNAPII-IPed samples as follow: First a nuclei isolation was performed as previously described. Purified nucleus were recovered in 300 μl of Nuclei Lysis Buffer (50 mM Tris-HCl pH8; 0.1% SDS; 10 mM EDTA; 100 μM PMSF; RNasin PROMEGA) and 5 cycles (30 s ON – 30 s OFF) of cell disruption applied with a refrigerated PicoRuptor. The samples were centrifuged for 10 min at 13000 g, 4°C and the supernatants were incubated overnight with 1/100 RNAPII or HIS antibodies (RNAP II AS11 1804, HIS AS20 4441, Agrisera) and 30 μl SureBeads (BioRad). Three washes were performed with ChIP Dilution Buffer as in DRIP assays before incubating the IP fraction with Proteinase K for 2 h at 55°C plus 15 min at 95°C in PK Buffer (100 mM Tris-HCl pH8; 50 mM NaCl; 10 mM EDTA; 4 mg/ml Proteinase K; RNasin PROMEGA). We added 2 U DNase I (ThermoFisher) and incubated it for 30 min at 37°C. After that, we continued with RNA purification as we describer before.

The 5′ RACE method was performed as as described previously (Llave et al., 2002). In brief, 2 μl of IP-RNA, or input RNA, were ligated to a RNA adapter (GeneRacer™ RNA, Thermofisher). Next, we used random primers for first strand cDNA synthesis. PCRs were performed with pri-miRNA specific reverse primers (Table S2). The amplification products were purified, cloned in pGEMT-easy vectors, and sequenced.

Small RNA sequencing of *hst-15* mutants was previously described (Cambiagno *et al*., 2021). Previously described AGO1-associated miRNAs datasets (Mi et al., 2008) were used to estimate loading of each miRNA. For this, we first calculated the fraction each miRNA represents to the total number of read in the input and IP samples. Then a ratio between the fractions in the IP/input was calculated to estimate the loading preference of each miRNA.

### Bioinformatics analysis

Chromosomal coordinates of *Arabidopsis thaliana* miRNAs were downloaded from the miRbase sequence database v22 (Kozomara et al., 2019). Hairpin precursor coordinates (annotated as “miRNA primary transcript”, pri-miRNA) and mature miRNAs coordinates were sorted by their biogenesis direction as indicated by (Moro *et al*., 2018) and each group was analyzed separately. The pri-miRNAs were scaled in order to exclude 5’ and 3’ arms to avoid noise signals when profiling plaNET-seq data. BTL, LTB, and LTBs pri-miRNAs were scaled from the miRNA-5p to the miRNA-3p chromosomal coordinates, and BTLs pri-miRNAs from the first DCL1 cut to the miRNA-3p genomic coordinates. Custom R scripts were written for this purpose and annotation files were manually curated. For the analysis of R-Loops formation over MIRNA loci, two coordinates windows were defined as follows: one from the TSS to the most up-stream DCL1 cleavage site, and the other from this site to the end of miRNA-3p. To define the first window, each pri-miRNA TSS was annotated de novo by combining information from different sources: PTSmiRNA database (You et al., 2017), TSS of Arabidopsis MIRNA primary transcripts reported by (Xie et al., 2005), mapped plaNET-seq reads (Kindgren *et al*., 2020), Arabidopsis thaliana ESTs and Full lenght cDNAs (Campbell et al., 2014), paired-end analysis of transcription start sites (Morton et al., 2014) and genome-wide TSS sequencing (Nielsen et al., 2019).

Samples from selected sequencing studies (Table S2) were downloaded from public repositories in bigWig format. The ssDRIP-seq mapped reads are available as forward (wR-loops, representing an R-loop formation containing ssDNA on the Watson strand and an DNA:RNA hybrid on the Crick strand) and reverse reads (cR-loops; (Xu *et al*., 2017)). In addition, reads from selected samples of plaNET-seq experiments were downloaded in SRA format and converted to fastq format using fasterq_dump (SRA-Toolkit, https://trace.ncbi.nlm.nih.gov/Traces/sra/sra.cgi?view=software).

Trimming, alignment to TAIR10 genome and post-processing of plaNET-seq reads were done as previously described (Kindgren *et al*., 2020; Leng *et al*., 2020), using the 01-Alignment_plaNET-Seq.sh and 02-Postprocessing_plaNET-Seq.R scripts available in the code repository: https://github.com/Maxim-Ivanov/Kindgren_et_al_2019. The script loadNETSeqBAM.R was modified in line 65 to obtain the genomic coverage in the 5’ nucleotide of mapped reads (mode = “start”) or the full coverage (mode = “whole_read”). In each case, genomic coverage was exported as strand-specific bigWig and bedGraph files using rtracklayer_1.42.2.

For the preparation of metagene plots of plaNET-seq data, the two biological replicates of each sample were merged using the bigWigMergePlus tool (https://github.com/c3g/kent/releases/tag/bigWigMergePlus_2.0.0). For some plots, strand-specific files were shown separately. The deepTools suite (Ramirez et al., 2016) was employed to draw metagene plots of plaNET-seq and ssDRIP-seq samples. ComputeMatrix tool was used in the scale-regions mode followed by plotProfile tool (parameters used are described in Table S3).

To calculate the ratio of co-transcriotional vs. post-transcriptinal processing each individual pri-miRNA, reads ending at DCL1 cleavage site (co-transcriptionally processed) and those expanding the site (unprocessed pri-miRNAs) were used (Figure 4A). For each independent experiment analyzed, the output files in bigWig format (plaNET-seq for co-transcriptional processing, and full coverage of re-mapped plaNET-seq reads for unprocessed pri-miRNAs) were used to calculate the scores for specific genomic regions (DCL1 cuts) using deepTools multiBigwigSummary in BED-file mode (Ramirez *et al*., 2016). In order to simplify the analysis, MIRNA loci with low plaNET-seq signal and with unclear processing mechanisms were excluded. The outputs of multiBigWigSummary were processed to obtain the ratios of pri-miRNAs in each condition and sorted by processing type in the R statistical programming environment (R Core Team, 2020), and graphics were produced with the ggplot package. Snapshots of the mapped plaNET-seq reads were constructed using the Integrative Genomics Viewer(Robinson et al., 2011).

### Probes preparation

For the detection of pri-miRNAs, we applied antisense DNA oligonucleotides labeled with digoxigenin at their 5’ ends and hybridizing to different segments of pri-miRNA163 and pri-miRNA156a. We designed probes targeting: an intron located downstream of the stem-loop structure (Intron), an exon (Exon), and two joined exons (Exon/Exon) as well as a loop sequence (Loop), a miRNA star (miRNA*) and a mature miRNA (miRNA) (check Table S1 for probe sequences). Next, we used Terminal Transferase Reaction to add an additional nucleotide conjugated with digoxigenin to the 3’ end of each probe according to the protocol delivered by Thermo-Fisher. The probes were incubated in reaction buffer (5 mM CoCl_2_, 400 U/reaction of Terminal Transferase (Sigma Merck) 0.1 mM DIG-11-dUTP (Sigma Merck), 0.1 mM dATP, 0.2 mM Alexa Fluor 488-5-dUTP (Thermo Fisher), probe final concentration: 10 pM, for 40 min. at 37° C.

RNA Stellaris probes were designed by using software: *Stellaris Probe Designer version 2.0* from Biosearch Technologies. The probes used were selected to target the intron sequence located downstream of the stem-loop structure of pri-miRNA156a and labeled with Quasar 570 or fluoresceine (6-FAM).

### Combined FISH and immunolocalization

pri-miRNAs were localized by applying fluorescence *in situ* hybridization (FISH) combined with the immunolocalization of digoxigenin attached to 5’ and 3’ ends of the probes. In our experiments FISH preceded the immunocytochemical methods. Prior to the assay, the cells were treated with PBS buffer containing 0.1 % Triton X-100 for cell membrane permeabilization. Next, the cells were hybridized with the probes in hybridization buffer (30% (v/v) formamide, 4× SSC (600 mM NaCl, 6 µM sodium citrate), 5× Denhardt’s solution (0.1% (g/v) ficoll 400, 0.1% (g/v) polyvinilpyrolidone, 0.1% (g/v) BSA), 1mM EDTA, and 50mM phosphate buffer) in a humified chamber for overnight at 26 °C. After washing, we applied the immunolabeling assay using primary mouse (Sigma Merck) or rabbit (Sigma Merck) anti-DIG antibodies (diluted 1:100) in 0.05% acetylated BSA in PBS for overnight at 10 °C. Subsequently, the cells were washed with PBS and subjected to incubation with goat anti-mouse or goat anti-rabbit secondary antibodies conjugated with Alexa Fluor 488 or Alexa Fluor 555 (Thermo Fisher) in 0.05% acetylated BSA in PBS for 2h at 37 °C. DNA was stained with Hoechst 33342 (Thermo Fisher) and mounted in ProLong Gold antifade reagent (Life Technologies).

### Immunodetection of proteins

Double immunodetection experiments were performed according to Bhat et al. (2020). The isolated nuclei were treated with PBS containing 0.1 % Triton X-100 and next incubated with primary antibodies in 0.05 % acetylated BSA in PBS for overnight at 10 °C. In our experiment we used antibodies targeting: SERRATE (Agrisera, diluted 1:100), HYL1 (Agrisera, diluted 1:200) and DCL-1 (Agrisera, diluted 1:100). For the localization of RNAPII we used antibodies recognizing RNAPII phosphorylated at serine 5 (Chromotek, diluted 1:200) and serine 2 (Chromotek, diluted 1:200). After washing with PBS, the slides were incubated with secondary goat anti-rabbit or goat anti-rat antibodies conjugated with Alexa Fluor 488 or Alexa 555 (Thermo Fisher, diluted 1:100) in PBS containing 0.01 % acetylated BSA at 37 °C for 2h. Next, the slides were stained for DNA detection with Hoechst 33342 (Thermo Fisher), and mounted in ProLong Gold antifade reagent (Life Technologies).

### Microscopic and correlation analysis

The results were registered with the Leica SP8 confocal microscope using lasers emitting light at wavelengths of 405, 488 and 561 nm with an optimized pinhole, long exposure time (200 kHz) and 63x (numerical aperture, 1.4). For the Leica confocal microscope Plan Apochromat DIC H an oil immersion lens was used. To minimize bleed-through between fluorescence channels, the low laser power (0.4–5% of maximum power) and single-channel collection were applied.

Correlation analyses were performed with the use of Pearson’s correlation coefficient, Spearman’s rank correlation and ICQ value by using free software *ImageJ* from the National Institute of Health in USA, and its plugin *coloc2* which is the analysis option of the expanded *ImageJ* version *Fiji*.

## Author contribution

L.G, I.T, T.G, Z.S-K, A.J and P.A.M conceived and designed the study. L.G, performed all IP experiments and validations; I.T. analyzed all the sequencing data; T.G. and A.K-M performed all microscopic experiments; D.A.C performed selected HST experiments; J.D.S designed and supervised microscopic experiments; P.A.M, A.J, Z.Sz-K and S.M supervised the work and secured project funding; L.G, I.T, T.G, Z.Sz-K, A.J and P.A.M wrote the manuscript with input from all authors.

## Acknowledgments

This work was supported by grants from ANPCyT (Agencia Nacional de Promoción Científica y Tecnológica, Argentina) and the Polish National Science Centre (UMO-2019/32/T/NZ1/00508, UMO-2016/23/N/NZ1/00010, UMO-2013/10/A/NZ1/00557). The authors also received financial support from the Initiative of Excellence–Research University (05/IDUB/2019/94) at Adam Mickiewicz University, Poznan, Poland. P.A.M. and D.A.C. are members of CONICET; L.G is a fellow of the same institution. We would like to thanks Federico D. Ariel for his valuable comments on this project and Dr. Pauline Julline (Institute of Plant Science, University of Bern, Switzerland) for sharing with us seeds of transgenic plants expressing mCherry-AGO1

## Footnotes

The authors declare no competing interest.

