## Supplemental information for "R-Loops between nascent pri-miRNAs and the encoding loci promote co-transcriptional processing of miRNAs in plants"

**Supplemental Figures**

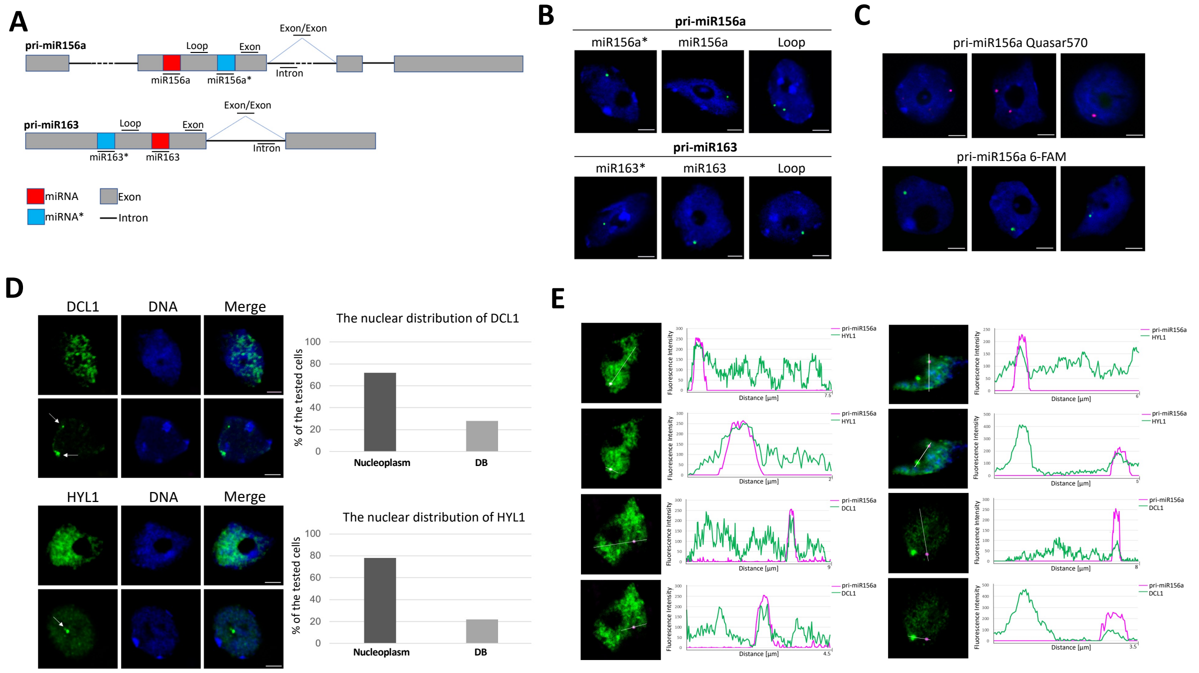


**Figure S1. Subcellular distribution and interactions of nascent pri-miRNAs and the miRNA biogenesis complex.**

(A) Schematic diagram of pri-miRNA163 and pri-miRNA156a in *A. thaliana*. Colors indicate structural elements of pri-miRNA: exon (grey bar), intron (dark line), mature miRNA (red bar) and miRNA star (blue bar). The probes used are depicted as green above each scheme.

(B) FISH of pri-miRNA156a and pri-miR163 (green) using mouse antibodies targeting digoxigenin in the nuclei isolated from wild-type plant cells. The probes hybridizing to the loops (Loop), miRNAs star (miRNA*), and mature miRNAs (miRNA).

(C) Fluorescence in situ hybridization (FISH) of pri-miR156a using RNA Stellaris probes. Representative images of cell nuclei showing the subnuclear localization of pri-miRNA156a using RNA Stellaris probes labeled with Quasar 570 (top) or fluorescein 6-FAM (bottom).

(D) Immunolocalization of DCL1 and HYL1 (green) in nuclei of wild type plant cells. Two types of distribution are shown: dispersed in the nucleoplasm (upper rows) and accumulated in D-bodies (lower rows). Percentage of nuclei with the dispersed distribution and the nuclei with visible D-bodies are shown in the right.

(E) Co-localization of pri-miRNA156a with HYL1 and DCL-1 in isolated nuclei of *A. thaliana* cells*.* FISH of pri-miRNA156a (magenta) combined with immunolabeling of HYL1 or DCL1.

Cells presenting a disperse or D-bodied localized DCL1 and HYL1 distribution are shown. On the right of each image is the fluorescence intensity plot along the white line draw in the microscopy picture. The fluorescence intensity of pri-miR156a is depicted as the magenta curve and HYL1/DCL1 as the green curve.

In all cases, the nuclei were stained with Hoechst (blue). Scale bar - 2.5 µm.

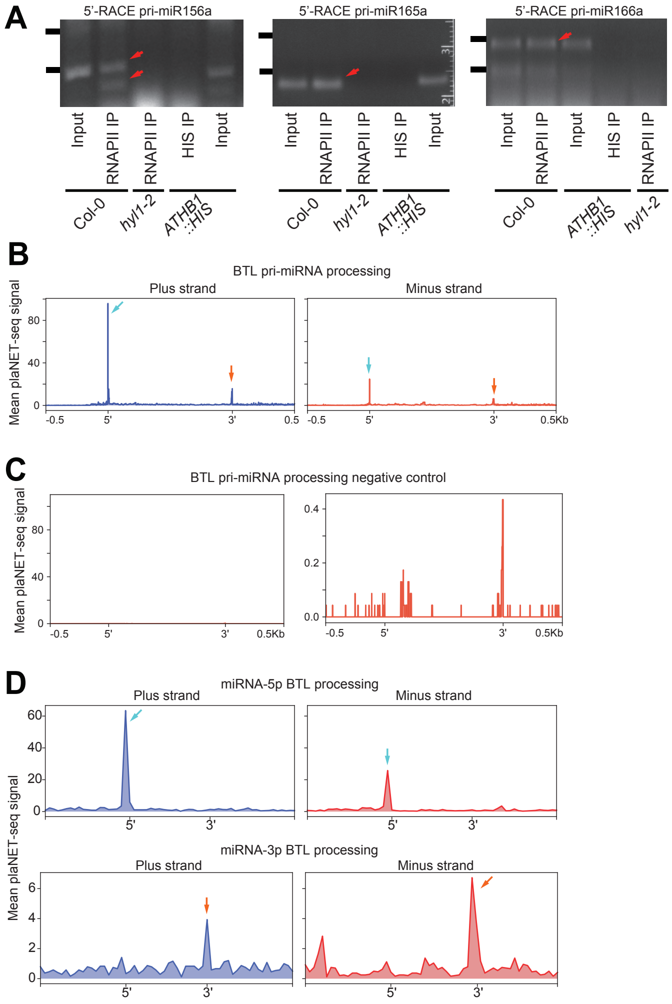


**Figure S2. Plant pri-miRNAs are processed co-transcriptionally.** (A) Amplification of pri-miRNA processing intermediates associated with RNAPII as detected by 5'RACE. Bands noted with red arrows were cloned and used to score processing intermediates. (B-D) Metagene analysis of nascent BTL pri-miRNAs processing intermediated associated to RNAPII as determined by scoring the 5'-end (B and C) or 3'-end (D and F) nucleotide of plaNET-seq reads. Pri-miRNAs were scaled from the beginning of miRNA-5p to the end of miRNA-3p (B and D) or using only the mature miRNAs sequences (C and F). Cyan and orange arrows indicated the processing fragments detected also marked in the schemes illustrated next to panels (B) and (D).

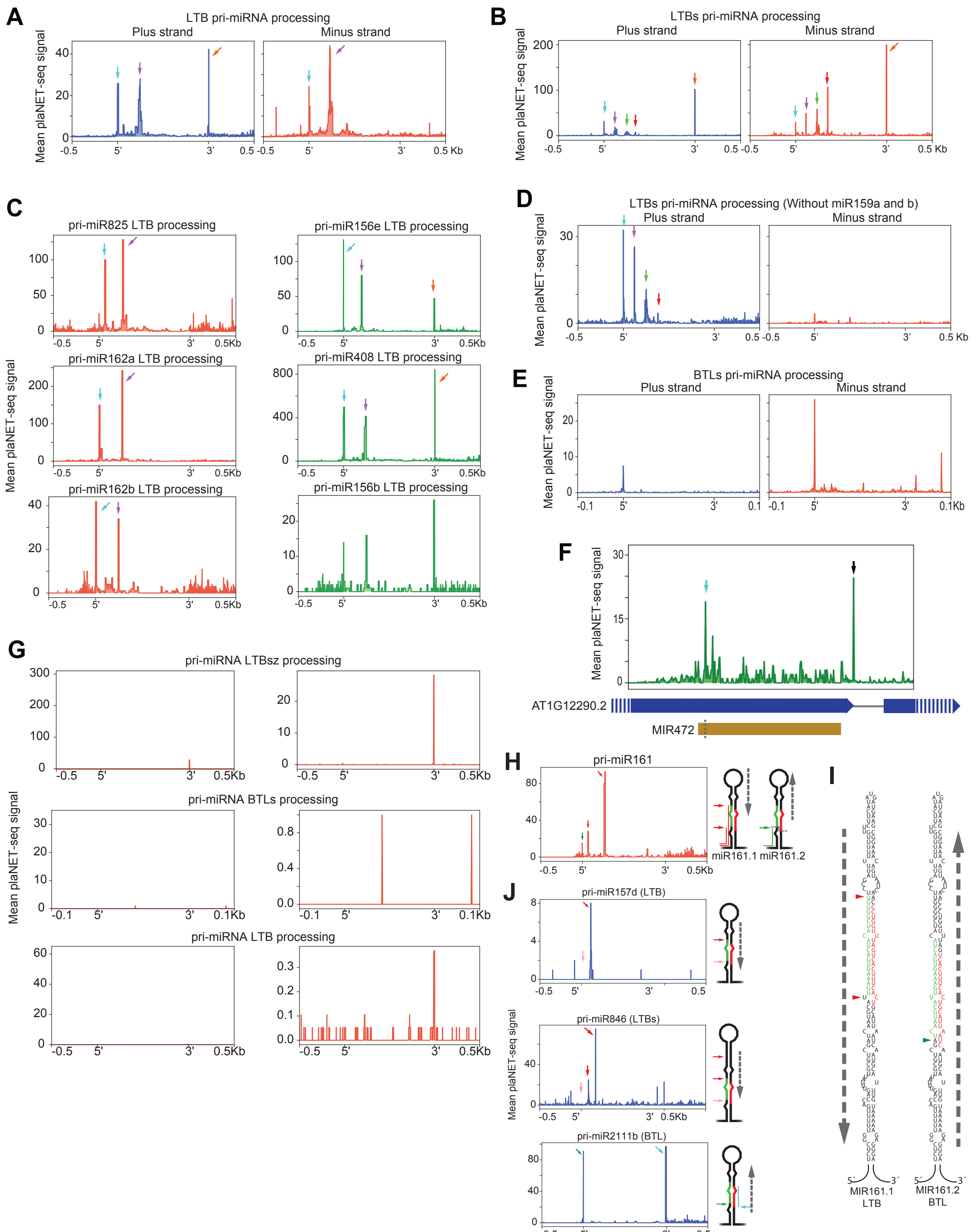

**Figure S3 Co-transcriptional processing of BTL pri-miRNAs involves a second nucleoplasmic processing step.** (A-G) Metagene analysis of nascent pri-miRNAs processing intermediated associated to RNAPII as determined by scoring the 3'-end nucleotide of plaNET-seq reads. Metagene analysis of loop to base pri-miRNAs (A), sequential loop-to-base pri-miRNAs (B), individual LTB loci (C), filtered sequential loop-to-base pri-miRNAs (D), sequential base-to-loop pri-miRNAs (E), and *MIR472* locus (F). In all cases color arrows correspond to positions indicated in Figure 3 and described in the result session. (F) The position of MIR472 within AT1G12290 is marked in yellow with the DCL1 cleavage site noted as a dashed line. AT1G12290.2 exons are noted with blue boxes. A peak mapping to the exon donor site is noted with a black arrow. (G) Metagene analysis of LTB, LTBs, and BTLs nascent pri-miRNAs processing intermediated in mock FLAG-IP negative controls samples. Left panels show an scale identical to the corresponding sample in Figure 3 while right panels show a zoom in.

(H) PlaNET-seq signal profile of miR161 which processing mechanisms was previous inferred BTL exclusive and now defined as dual BTL-LTB. Pri-miRNA was scaled from the beginning of miRNA-5p to the end of miRNA-3p. Color arrows indicate DCL1 processing site marked in the adjacent schemes and in panel (I).

(I) Predicted secondary structures of pri-miR161.1, and pri-miR161.2. Color arrow heads indicate the position of plaNETseq peaks displayed in panel (H). The sequence corresponding to the mature miRNA-5p is displayed in green and the miRNA-3p in red. Determined processing direction is noted with dashed arrows next to the structures.

(J) Identification of co-transcriptional processing mechanisms of miR157d, miR2111b, and miR846 based on plaNET-seq signal profiles. Pri-miRNAs were scaled from the beginning of miRNA-5p to the end of miRNA-3p. Color arrows indicate DCL1 processing site marked in the adjacent schemes. Light red arrows indicate expected cleavage sites not detected probably due to low coverage of the analyzed loci. Cyan arrow in pri-miR2111b indicates a peak corresponding to retention of the mature miRNA-3p in the processing complex.

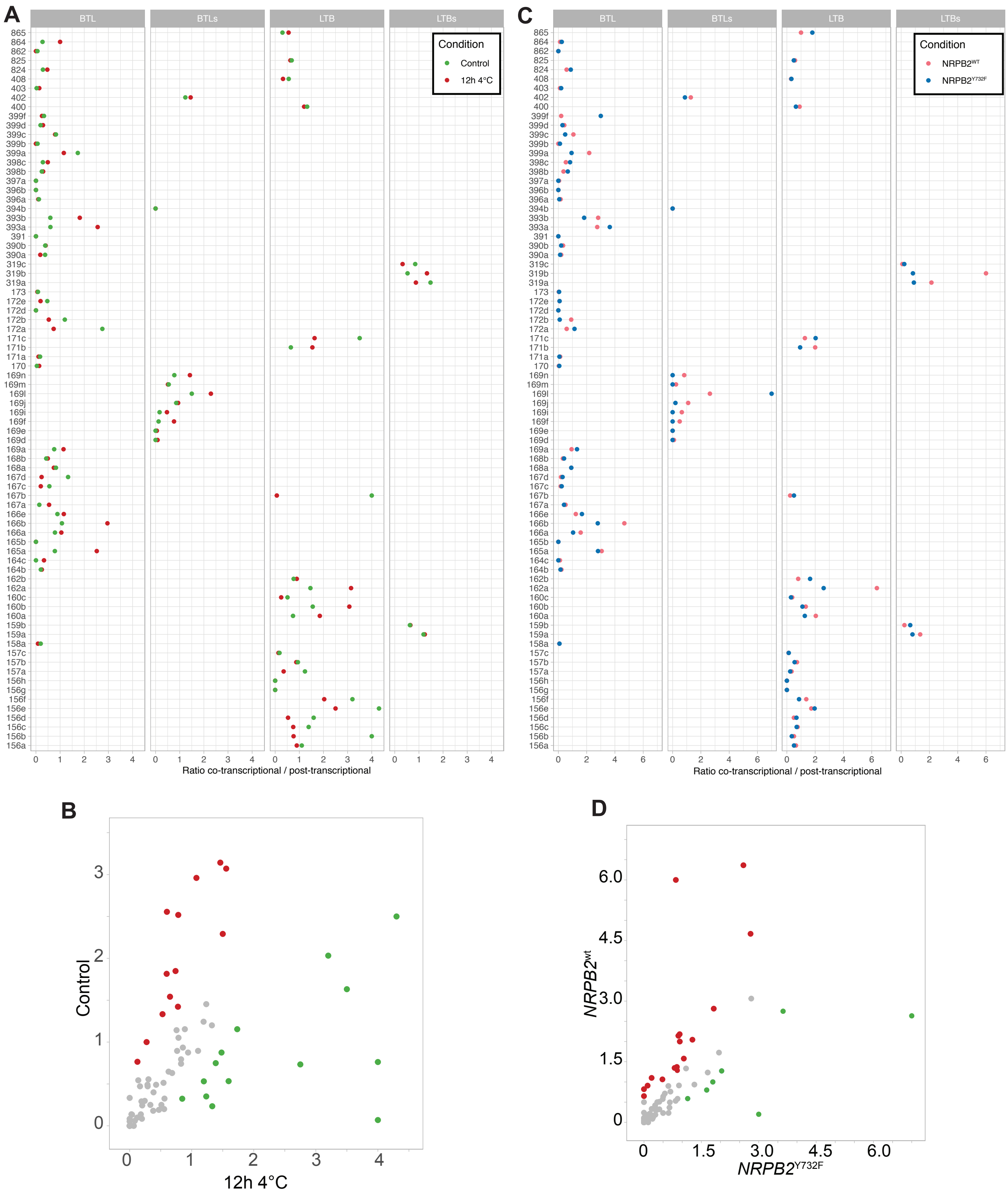


**Figure S4. Co-transcriptional miRNA processing ratios are variable in different conditions.**

(A and C) Co-transcriptional processing ratios corresponding to all analyzed miRNAs in plaNET-seq experiments performed in control plants or in plants incubated 12 h at 4 °C (A) or transformed with a WT or Y732F mutant versions of NRPB2 (B) and split by processing mechanism.

(B and D) Scatter plot representations of the co-transcriptional processing ratios in the same samples described above. Red and green dots show those pri-miRNAs with higher or lower co-transcriptionally processing ratios in control conditions.

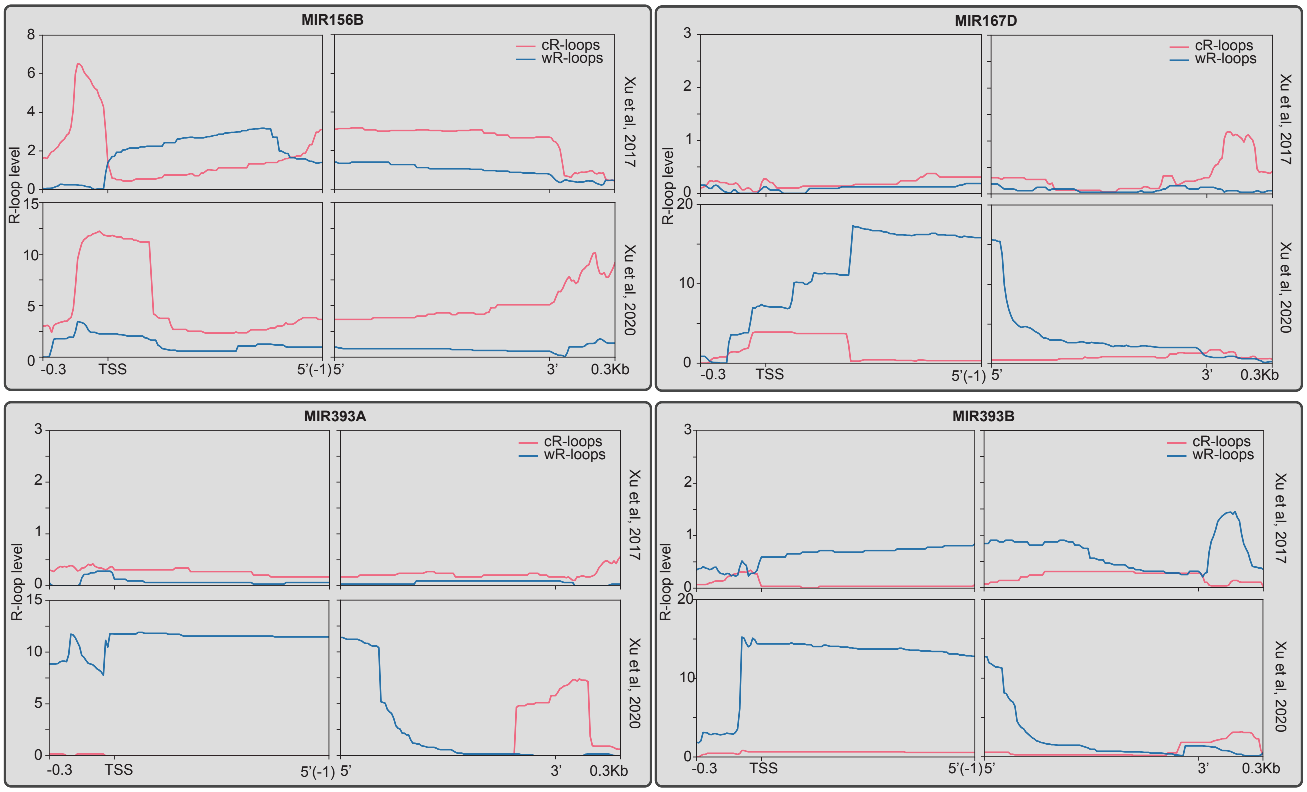


**Figure S5. R-Loops at the 5' end of miRNA loci promotes co-transcriptional processing.** R-loop formation over selected *MIRNA* loci either in the Watson (blue) or Crick (red) strands in samples prepared in studies using different developmental stages.

**Table S1.** List of primers used in this study.

| Gene (AGI) | Sequences (5´ - 3´) | Assay |
|---|---|---|
| pri-miR165a | CCATCATCACCATTCACCAACC | qPCR |
| | CCTCAACTGAAATAGCTTAACCC | qPCR |
| | GTTGTCTGGATCGAGGATATTATAG | qPCR |
| | GTCCGAGGATACTCTCTATGATC | qPCR |
| | ACATGTTATTGCCTCTGATCACC | qPCR |

| | | |
|---|---|---|
| | GCAAGAAAGATTCAAAGTCATCAC | qPCR - 5'RACE |
| pri-miR166b | GGATCTGTTGGGGGACGAAC | qPCR |
| | CCTCAAAAGAAAAATCCCTC | qPCR |
| | GGCTCGAGGACTCTTATTC | qPCR |
| | CCGACGACACTAAAACCC | qPCR |
| | CAATTATCACTCCCTCACAATCC | qPCR |
| | CACATGGATTCATAGATAGAAACC | qPCR |
| pri-miR168a | CATATCATAAACCTCATTTCCCCA | qPCR |
| | CGAGCCCGATGGTGAGACTC | qPCR |
| | GGAACCAATTCGGCTGACAC | qPCR |
| | GGGATCCAATCCCTGCTCAC | qPCR |
| | TCCTCGAGGTGTAAAAAAACTCG | qPCR |
| | GAGAAGCGTAGAAATCTTCCAG | qPCR |
| pri-miR159a | TGGTAGAGCTCCTTAAAGTT | qPCR |
| | TAAAGCTCCTGAGATATGCA | qPCR |
| | GGTCTTTACAGTTTGCTTATG | qPCR |
| | TAAAGCTCCTGAGATATGCA | qPCR |
| | CGACTCTCTATCTATCATTTCTTCC | qPCR |
| | GAAAGTTTTGATGAGAACGTGG | qPCR |
| pri-miR156a | GTGAGCACGCAAGAGAAGCAAG | qPCR |
| | AAGAAACTGACAGAAGAGAGTGAG | qPCR |
| | GAGAACGAAGACAGGCCAAAG | qPCR |
| | AAGAAACTGACAGAAGAGAGTGAG | qPCR |
| | ATCTTGTAGATCTCTGAAGTTGG | 5'RACE |
| pri-miR408 | GACAGGGAACAAGCAGAGCATGG | qPCR |
| | GAGACAAAACAGAGTCGTTTAATG | qPCR |
| | GAACTAACTCAAAGGAACTGGC | qPCR |
| | GAGACAAAACAGAGTCGTTTAATG | qPCR |
| | CTCTCTCATTACCGCTTTGTCTTC | qPCR |
| | CATGCTCTGCTTGTTCCCTGT | qPCR |

| | | |
|---|---|---|
| pri-miR402 | GGTTCATCGGAAGGAGTTAGC | qPCR |
| | GCAACTCAAACTTATCTACCACG | qPCR |
| | GGTTCATCGGAAGGAGTTAGC | qPCR |
| | GAATCTGCTGTTGGAAATTGAGGC | qPCR |
| | CATGAAAATATGGGTTAACAAACAAAGG | qPCR |
| pri-miR160b | CTCTGGTTCATGTTTTCCCC | qPCR |
| | GAACCCTTAAATATGATTGGTGG | qPCR |
| | CTCTGGTTCATGTTTTCCCC | qPCR |
| | TGCTTGACTACTCTGTACG | qPCR |
| pri-miR166a | TAACAAGGGTTCATTCACTGG | 5'RACE |
| | CTGGCTCGCTCTATTCATGTTGG | qPCR |
| | GACGCTAAAACCCTAATCAAATC | qPCR |
| | GGGACGAACATAGAAAGAGAGAG | qPCR |
| | GCCCCTTTTTCTTTTCAGTCG | qPCR |
| pri-miR164b | CAGACAAATCATACCCCCAAGG | qPCR |
| | CACACCTTCATCATTCTCTCCG | qPCR |
| | CCACAAATGCGTGTATATATGC | qPCR |
| | CTAACTCATCCATATCATCACACTC | qPCR |
| MIR319b | TCTAGCACGCACAGAGAGG | qPCR |
| | CCATGAGTGGACCGAAGAAAGC | qPCR |
| MIR171a | TTTCCTTTGATATCCGCACTTTAAG | qPCR |
| | GATATTGGCGCGGCTCAATC | qPCR |
| APOLO | CGGGGACATCCGATAAAATTGG | qPCR |
| | CGATTTGTGCGTGTCATCCTTG | qPCR |
| U6 | CGGGGACATCCGATAAAATTGG | qPCR |
| | CGATTTGTGCGTGTCATCCTTG | qPCR |
| pri-miR163 Intron | GTTAATGTTAGTAGTTAAAAAGGATTAGTG | Probe |
| pri-miR163 Exon | TACGTTATCTCTTTTCATCAATTAAACC | Probe |
| pri-miR163 Exon/Exon | CAAAAAATTTCCGTTATCTCTTTT | Probe |
| pri-miR163 Loop | GGAACTCCAGCACTTTAGTATCATC | Probe |
| miR163* | GAAGAGGTTGGAACTCGATTT | Probe |

| | | |
|---|---|---|
| miR163 | ATCGAAGTTCCAAGTCCTCTTCAA | Probe |
| pri-miR156a Intron | GCTAATCTCACTTAACCACGCG | Probe |
| pri-miR156a Exon | AGAGAGATTGAGACATAGAGAACG | Probe |
| pri-miR156aExon/Exon | ACCCCCTTACCTTAATATGG | Probe |
| pri-miR156a Loop | TGAGCACGCAAGAGAAGCAAGT | Probe |
| miR156a* | TGACAGAAGAGAGTGAGCA | Probe |
| miR156a | GTGCTCACTCTCTTCTGTCA | Probe |

**Table S2. IDs of the datasets used in this study.**

| **Study** | **Ref.** | **Series Accession** | **Samples analyzed** | **Description** |
|---|---|---|---|---|
| Transient genome-wide adaptations of nascent RNAPII transcription triggered by low temperature | {Kindgren, 2020 #56} | GSE131733 | GSM3814845<br>GSM3814846<br>GSM3814849<br>GSM3814850 | Plant native elongating transcripts sequencing (plaNET-Seq) was used to determine the genomic positions of transcriptionally engaged RNAPII in Col-0 seedlings which were exposed to cold (4 °C) for 0h, 3h or 12h. |
| Organismal Benefits of Transcription Speed Control at Gene Boundaries | {Leng, 2020 #57} | GSE133143 | GSM3900879<br>GSM3900880<br>GSM3900881<br>GSM3900882 | Plant native elongating transcripts sequencing (plaNET-Seq) was used to determine the genomic positions of transcriptionally engaged RNAPII in wild type and nrpb2-Y732F seedlings. |
| R-loop landscapes in Arabidopsis | {Xu, 2020 #71} | GSE116232 | GSM3214368<br>GSM3214369<br>GSM3214344<br>GSM3214345<br>GSM3214346<br>GSM3214347<br>GSM3214348<br>GSM3214349 | ssDRIP-seq in Arabidopsis to profile genome-wide R-loop levels providing a first-hand R-loop atlas during Arabidopsis development and in response to various environmental factors. |

| | | | | |
|---|---|---|---|---|
| | | | GSM3214382 GSM3214383 GSM3214328 GSM3214329 | |
| Genome-wide maps of R-loops in Arabidopsis | {Xu, 2017 #42} | GSE95765 | GSM2525600 | ssDRIP-seq in 12-day-old Arabidopsis seedlings for genome-wide identification of R-loops. |

**Table S3. Deeptools parameters used to scale different regions for metagen analyses using ComputeMatrix and plotProfile tools.**

| **Mode** | **Genome regions** | **Score files (.bw) from:** | **--regionBodyLength** | **--upstream** | **--downstream** | **--binSize** | **Figure** |
|---|---|---|---|---|---|---|---|
| scale-regions | pri-miRNAs (previously scaled and sorted by processing type) | PlaNET-seq (5’ nt) | 1000 bp | 500 bp | 500 bp | 1 | 1B |
| scale-regions | pri-miRNAs (previously scaled and sorted by processing type) | PlaNET-seq (3’ nt) | 1000 bp | 500 bp | 500 bp | 1 | 1D, S1B, 2A-2D, S2A-D, S2F, 5A-E, S5D |
| scale-regions | Mature miRNAs (miRNA-3p or miRNA-5p) | plaNET-seq (5’ nt) | 20 bp | 20 bp | 20 bp | 1 | 1C |
| scale-regions | Mature miRNAs (miRNA-3p or miRNA-5p) | plaNET-seq (3’ nt) | 20 bp | 20 bp | 20 bp | 1 | 1F, S1C |
| scale-regions | Mature miRNA-3p | plaNET-seq (3’ nt and whole_read) | 40 bp | 20 bp | 20 bp | 1 | 2E |

| | | | | | | | |
|---|---|---|---|---|---|---|---|
| scale-regions | pri-miRNAs (previously scaled) | PlaNET-seq (3’ nt) | 200 bp | 100 bp | 100 bp | 1 | 2G, S2E |
| scale-regions | pri-miRNAs | PlaNET-seq (3’ nt) | 1000 bp | 500 bp | 500 bp | 1 | 6G, 6H |
| scale-regions | Two windows:<br>1) TSS to most up-stream DCL1 cleavage site<br>2) pri-miRNA (previously scaled) | ssDRIP-seq | 1) 1000 bp<br>2) 1000 bp | 1) 300 bp<br>2) 0 bp | 1) 0 bp<br>2) 300 bp | Default: 10 | 6C, 6E, S6 |
| scale-regions | Coordinates regions including polycistronic miRNA duets or miRNAs encoded within protein coding genes | ssDRIP-seq | 1000 | 300 bp | 300 bp | Default: 10 | 6G, 6H |
